# RNAchat: Integrating machine learning algorithms to identify metapathways based on clinical and multi-omics data

**DOI:** 10.1101/2025.04.02.646761

**Authors:** Mingcan Tang

## Abstract

**Background:** Understanding the global interactions among cellular pathways and types is essential for unraveling the complex biological mechanisms that underlie various diseases. While many existing studies have primarily focused on omics data, they often fail to integrate clinical heterogeneity, which is crucial for a comprehensive understanding of disease biology. Although significant progress has been made in researching crosstalks between inter-pathways and inter-cells, and tools like CellChat have emerged to analyze these interactions, they have notable limitations. These include the assumption that mRNA expression directly reflects protein activity, a heavy reliance on pre-established ligand-receptor interactions, and an inability to connect “chats” with clinical phenotypes. Furthermore, existing methods have limitations in distinguishing between paracrine signaling (interactions between different cell types) and autocrine signaling (self-signaling within the same cell type). These limitations hinder the ability to translate molecular insights into phenotypic outcomes, thereby limiting clinical applicability. To provide another solution for these challenges, we introduce RNAchat, a novel approach that utilizes machine learning techniques to identify metapathways for crosstalks among pathways at not only the bulk level but also cell types at the single-cell level, incorporating both clinical and multi-omics data.

**Results:** RNAchat provides another solution by providing an interactive, reproducible platform designed to analyse metapathways between inter-pathways and inter-cell types using multi-omics level data in a clinical context. This tool enables researchers to integrate diverse datasets, conduct exploratory analyses, and identify cooperated pathways. The platform facilitates hypothesis generation and produces intuitive visualizations to support further investigation.

As a proof of concept, we applied RNAchat to connect omics data to pain, drug resistance in rheumatoid arthritis (RA), disease severity to RA and Heart Failure (HF). We discovered cooperated molecular pathways associated with different treatments using SHAP (Shapley Additive Explanations) values. Then, we demonstrated how to use it to find cellular interactions at single cell level. Last but not the least, we further extended crosstalk exploration between host and parasites and found potential novel important pathway interaction not mentioned by previous studies.

**Conclusion:** We introduce RNAchat, a computational platform designed to identify pathway communications with clinical and multi-omics data in real-time. This tool supports both user-generated and publicly available datasets, offering a robust solution and enhancing our understanding of complex diseases such as RA and HF. The platform is available at https://github.com/tangmingcan/RNAchat.

## BACKGROUND

Autoimmune and autoinflammatory diseases incorporate a heterogeneous group of chronic diseases with substantial morbidity and mortality, thereby posing significant health challenges. While there have been substantial advances in the treatment of RA and related rheumatic immune-mediated conditions, based on improved understanding of the underlying disease pathogenesis, there remains significant unmet clinical needs. Treatment responses remain unpredictable and inadequate response is common; some patients may respond to drugs with one mode of action, while others may respond to another drug or not at all ^1^. Furthermore, although recently with the development of machine learning, signatures were found by many previous studies connecting to drug resistance/pain in RA, it still lacks a solid theory to inter-connect patients’ clinical heterogeneity and multi-omic evidence with a generalised drug-independent treatment outcome ^2–4^.

RNAcare^5^ was developed based on tools of DEG analysis such as Phantasus^6^, CFViSA^7^, AmiC^8^, using Django, Regularisation Regression^9,10^ and distributed framework for accelerating the analysis process. It provides insightful results in terms of pain, fatigue and drug response for RA treatment. However, it didn’t consider interaction between metagenes; RNAcompare is the development of RNAcare, focusing on addressing the issues of heterogeneous data and suggesting its own solution for tissue-cross, omic-cross, batch-cross, tech-cross and disease-cross data. But it still has limitations: It found crosstalk/synergy effects passively by SHAP dependence plot, and it limits to only two metagenes; used Pearson Correlation matrix for variable selection while potentially retaining overlapped metagenes with same biologic function; didn’t consider crosstalk among pathways/cell types in a single cell background.

At the same time, softwares such as CellChat^11^ have appeared to analyze these interactions at the single cell level. Still, they have notable limitations. These include the assumption that mRNA expression directly reflects protein activity, a heavy reliance on pre-established ligand-receptor interactions, and an inability to infer/prioritise direct functional effects based on clinical outcome. Furthermore, existing methods struggle to distinguish between paracrine signaling (interactions between different cell types) and autocrine signaling (self-signaling within the same cell type) at single cell level.

MetaPathways^12^ is a bioinformatics software designed for functional annotation and metabolic pathway reconstruction from metagenomics and metatranscriptomic data. It is particularly useful for analyzing microbial communities and understanding their metabolic capabilities. However, it only focuses on metabolic and biochemical pathway interactions among microbes; still an unsupervised in nature, which means it lacks a pre-defined functional target.

To address these complexities, we introduce RNAchat. This development of RNAcompare offers an automated pipeline that seamlessly integrates clinical and multi-omics data for clinicians, facilitating the identification of metapathways across inter-pathways/inter-cells through advanced machine learning algorithms. By enabling comprehensive, multi-levels analyses, RNAchat aims to enhance our understanding of disease pathogenesis and inform the development of more effective, personalized treatment strategies.

## IMPLEMENTATION

The overall aim of RNAchat is to provide a web-based tool simplifying the identification of interactions among pathways based on multi-omics and clinical data and showing the analysis workflow in tabs as outlined in Figure 1 which will be described later in more detail. RNAchat was developed in Python, utilizing several open-source packages. The webserver was built using the Django framework, adhering to the FAIR^13^ (Findable, Accessible, Interoperable, and Reusable) principle. The platform employs the Plotly graphics system for generating interactive visualisations on the fly. The platform can be installed locally from https://github.com/tangmingcan/RNAchat.

**Figure 1.**
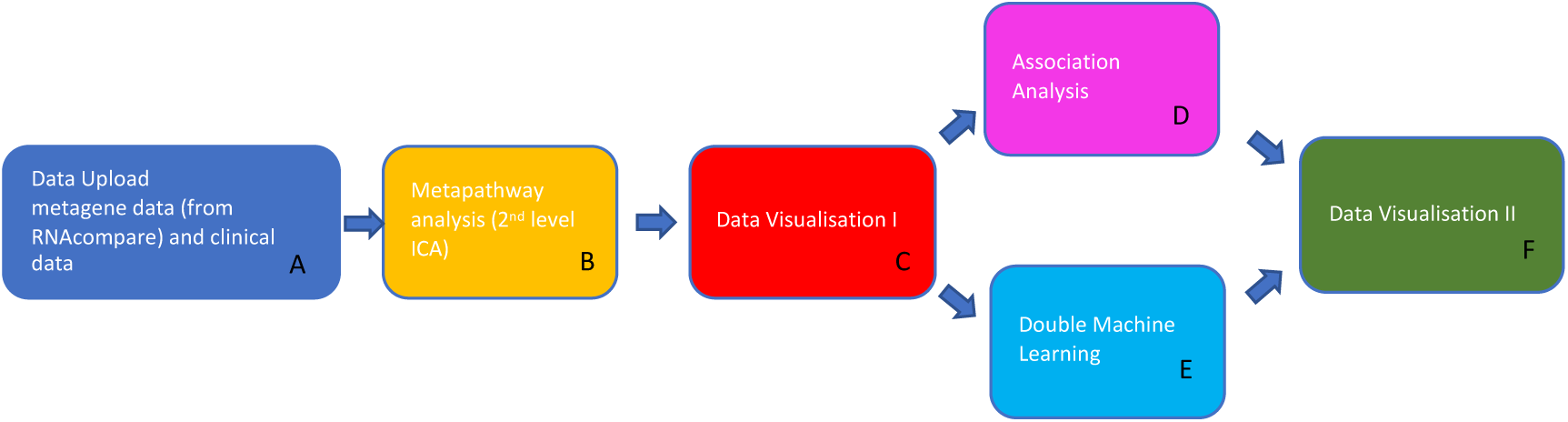
-RNAchat work flow. (A) In the first tab, user can upload processed metagene data from RNAcompare and clinical data; (B) Metapathway analysis where data was processed by Independent Component Analysis (ICA) again. (C) Data Visualisation I for the 2^nd^ level ICA processed data to check relationships and expression level of metapathways; (D) Association Analysis based on metapathways; (E) Double Machine Learning (DML) analysis based on metapathways; (F) Data Visualisation II for cooperated metagenes in the metapathways and cooperated cell types (only for single cell data).

To enable several users working on the system, Django is used to allow group and authentication management. System administrators can easily assign group roles to specific users, enabling them roles to only view and operate the permitted data.

### Datasets

We introduced the following RA, HF, single cell and malaria datasets (Table 2) – Optimal management of RA patients requiring Biologic Therapy (ORBIT^14^), Pathobiology of Early Arthritis Cohort (PEAC^15^), RA-MAP^16^ as part of the IMID-Bio-UK consortium, GSE135055^17^, RV infection scRNA-seq^18^, GSE288146 ^19^ and GSE289197 ^20^.

**Table 1.** Overview of datasets used in RNAchat. For RV infection scRNA-seq data, we put cell types in the clinical data.

| Cohort name | Number of participants | Data type | tissue type | #clinical feature |
| --- | --- | --- | --- | --- |
| PEAC1 | 49 | Bulk RNA-seq<br>(harmonised with ORBIT) | Whole Blood | 7 |
| PEAC2 | 67 | Bulk RNA-seq | Whole Blood | 1 |
|  | 53 | Bulk RNA-seq | Synovium | 1 |
| ORBIT | 139 | Bulk RNA-seq<br>(harmonised with PEAC1) | Blood | 16 |
| RA-MAP | 63 | Relative fluorescence<br>units(RFU) | Plasma | 1 |
|  | 34 | Microarray | Whole Blood | 1 |
| GSE135055 | 18 | Bulk RNA-seq-FPKM | Left ventricular heart<br>tissue | 2 |
| RV infection<br>scRNA-seq | 4 batches | Single cell RNA-seq | primary human<br>bronchial organoids | 1* |
| GSE288146 | 2 batches | Single cell RNA-seq | Plasmodium | 0 |
| GSE289197 | 79 | Bulk RNA-seq | Whole Blood | 1 |

**Table 2.** Metapathway analysis.

| Metagene 2<sup>nd</sup> level | Metagene 1<sup>st</sup> level | Enrichment result | Explanation |
| --- | --- | --- | --- |
| Metapathway_2(All) | 16 | REACTOME_RHO_GTPASE_CYCLE | I IFN |
|  | 14 | REACTOME_MHC_CLASS_II_ANTIGEN_PRESENTATION<br>KEGG_ALLOGRAFT_REJECTION | II IFN |
|  | 7 | REACTOME_TOLL_LIKE_RECEPTOR_CASCADES<br>REACTOME_SIGNALING_BY_INTERLEUKINS |  |
|  | 17 | REACTOME_CLASS_I_MHC_MEDIATED_ANTIGEN_PROCESSING_PRESENTATION | I IFN |
|  | 8 | REACTOME_RHO_GTPASE_CYCLE<br>WP_B_CELL_RECEPTOR_SIGNALING | I IFN |
|  | 0 | REACTOME_RESPONSE_TO_ELEVATED_PLATELET_CYTOSOLIC_CA2<br>REACTOME_FORMATION_OF_FIBRIN_CLOT_CLOTTING_CASCADE |  |
| Metapathway_6(Anti-TNF) | 6 | REACTOME_NONSENSE_MEDIATED_DECAY_NMD | I IFN |
|  | 17 | REACTOME_CLASS_I_MHC_MEDIATED_ANTIGEN_PROCESSING_PRESENTATION | I IFN, II IFN |
|  | 15 | REACTOME_EUKARYOTIC_TRANSLATION_ELONGATION<br>REACTOME_NONSENSE_MEDIATED_DECAY_NMD_INDEPENDENT_OF_THE_EXON_JUNCTION_COMPLEX_EJC | I IFN |
|  | 8 | REACTOME_RHO_GTPASE_CYCLE<br>WP_B_CELL_RECEPTOR_SIGNALING | I IFN |
| Metapathway_3(Rituximab) | 16 | REACTOME_RHO_GTPASE_CYCLE | I IFN |
|  | 14 | REACTOME_MHC_CLASS_II_ANTIGEN_PRESENTATION<br>KEGG_ALLOGRAFT_REJECTION | II IFN |
|  | 7 | REACTOME_TOLL_LIKE_RECEPTOR_CASCADES<br>REACTOME_SIGNALING_BY_INTERLEUKINS |  |
|  | 8 | REACTOME_RHO_GTPASE_CYCLE<br>WP_B_CELL_RECEPTOR_SIGNALING | I IFN |
|  | 17 | REACTOME_CLASS_I_MHC_MEDIATED_ANTIGEN_PROCESSING_PRESENTATION | I IFN, II IFN |
|  | 2 | REACTOME_L1CAM_INTERACTIONS<br>BIOCARTA_AHSP_PATHWAY | II IFN |

The Pathobiology of Early Arthritis Cohort (PEAC) was established with the aim to create an extensively phenotyped cohort of patients with early inflammatory arthritis, including RA, linked to detailed pathobiological data. The clinical and transcriptomic data can be found on EBI ArrayExpress with accession E-MTAB-6141. When analysing with ORBIT, we used processed PEAC1, downloaded directly from RNAcare after its own qualification control; When analysing based on two tissues, we used the original PEAC from the source mentioned.

ORBIT was a study comparing Rituximab to anti-TNF treatments. Blood was taken from patients before drug treatment. RNA-Seq data are being submitted to Array Express as pseudo-anonymized data. It’s likely that patients had previously been on csDMARDs such as Methotrexate, which had been stopped due to either lack of response or toxicity effects.

The RA-MAP Consortium is a UK industry-academic collaboration to investigate clinical and biological predictors of disease outcome and treatment response in RA, using deep clinical and multi-omic phenotyping. The raw data were retrieved from accessions GSE97810 and GSE97948 on ArrayExpress. The clinical data can be found here: https://doi.org/10.6084/m9.figshare.c.5491611.v1

GSE135055 was a study about Heart Failure (HF), collected from left ventricular heart tissue from 21 HF patients and 9 healthy donors as study cohort and generated multi-level transcriptomic data. We just used 18 patients with Dilated Cardiomyopathy (DCM), a condition where the heart’s left ventricle (the main pumping chamber) becomes enlarged and weakened, reducing its ability to pump blood effectively.

RV infection scRNA-seq data about rhinovirus infection in primary human bronchial organoids comprises four conditions: exposure to cigarette-smoke extract (CSE), rhinovirus (RV) infection, the combination of rhinovirus and cigarette smoke (RVCSE), and a control condition (mock). Although rhinovirus infection has been investigated^18^, the goal of our study was to further probe cellular communication to viral infection from each airway epithelial cell type in the presence or absence of a common environmental insult that is known to impact the outcome of rhinovirus infection: cigarette smoke. The data is available on Dryad^21^.

Malaria pathogenesis, encompassing parasite invasion, egress, and antigenic variation, relies on the coordinated activity of numerous proteins, yet their molecular regulatory mechanisms remain poorly understood. GSE288146 was an experiment for the intrinsic mechanism of PfAP2-V regulation in P. falciparum by scRNA-seq analysis. Here we just used the baseline data.

GSE289197 is transcript data detailing human transcription in severe malaria cases and uncomplicated malaria controls in a Mali study. Severe malaria cases were either cerebral malaria, severe malarial anemia, or concurrent cerebral malaria and severe malarial anemia disease.

### Clinical data used from the cohorts

RA disease activity was assessed using the validated DAS (Disease Activity Score using 28 joint counts) score, which was calculated from the original recorded 28 joint counts plus a blood marker of inflammation, typically the erythrocyte sedimentation rate (ESR) or the C-reactive protein (CRP) level^22^; DAS score in RA-MAP is standardised; Pain VAS^23^ is short for pain Visual Analog Scale (VAS) for self-reported pain, which is a unidimensional measure of general pain intensity, used to measure patients’ current pain level. The pain VAS is not specific to RA and has been widely used in a range of patient populations, including those with other rheumatic diseases, patients with chronic pain, cancer, or even cases with allergic rhinitis^24^, which provides an possibility for integration of different diseases for our later discussion.

HYNA (New York Heart Association^25^) is a system used to classify the severity of HF based on a patient’s symptoms and physical activity limitations. It has four functional classes: Class I: No symptoms and no limitations in ordinary physical activity; Class II: Mild symptoms and slight limitation during ordinary activities; Class III: Significant limitation in activity due to symptoms; comfortable only at rest; Class IV: Severe limitations; symptoms present even at rest.

To combine the disease severity of both diseases, we min-maximised each severity index (DAS in PEAC+ORBIT and HYNA in GSE135055) to 0-1 respectively, and then multiplied by 100. The second step is not necessary.

For RV infection scRNA-seq data, we put cell type of each record in the clinical data. All this clinical information is stored on csv files and, after curation, loaded into RNAchat.

For GSE289197, the number of parasitemia in the whole blood was used to measure the infection.

#### A - Data upload

The user has the option to import preprocessed data from RNAcompare, or upload their own ICA pre-processed data on the fly combined with clinical data which may include cell type information for scRNA analysis. Clinical data are uploaded as a table.

#### B - Metapathway Analysis

Based on the first-level independent component analysis (ICA^26^) processed data, we perform a second-level ICA on the metagenes. We hypothesize that the pathways represented by metagenes interact with one another, forming higher-order structures referred to as metapathways. Following the second-level ICA, we can extract the components of these metapathways, each composed of multiple metagenes. In the case of single-cell RNA sequencing (scRNA-seq) data, individual cell types are treated as specialized metagenes during the decomposition process. Consequently, the resulting metapathways may incorporate cell type information, reflecting ICA-derived correlations between cell types and pathways within the same metapathway. If a metapathway contains only a single cell type, it is indicative of autocrine signaling; if multiple cell types are present, it suggests paracrine signaling. Conversely, metapathways devoid of cell type representation correspond to generalized pathway interactions that are not cell type-specific. Metapathways can be directly downloaded after processing.

#### C - Data Visualisation I

In this section, users will perceive the relationships of metapathways. We provide 2 ways: Pearson Correlation Matrix^27^ and matrixplot scaled by metapathways/batches. When uses want to compare the expression level of different metapathways within the same tissue and omics level, the option of ‘scaled by metapathways’ is recommended; otherwise, “scaled by batches” is recommended.

#### D - Associatioin Analysis

Here, the user has the option to associate selected clinical data parameters with metapathway data in order to study phenotypes based on part or all of datasets. We provide Random Forests^28^, XGBoost^29^ and LightGBM^30^ with its corresponding parameters for basic tuning. The data will be split into training and test (20%) datasets by default. AUC-ROC^31^ or MSE^32^ will be shown according to different cases. This module will provide SHAP^33^ feature importance plot and SHAP dependence plot.

#### E - Double Machine Learning (DML) Analysis

We used DML introduced from RNAcompare to bypass data hamalisation for comparsion between different batches/tissues/omics levels. Causal Forests^34^ is recommended because it can estimate conditional treatment heterogeneity between batches, which can allow us to explore whether the most contributing factor (hub metapathways) in a given condition is the most important compared to protective mechanisms. Notably, the most significant contributor identified by Association analysis may differ from feature importance ranks obtained using Causal Forests.

#### F – Data Visualisation II

In this section, user can use network plot to visualise the interactions among metagenes/cell types for finding the most frequent/important metagenes/cell types.

## Results

Here we present the first tool that allows users without bioinformatics skill to integrate clinical and multi-omics data, establish an automated pipeline, and look for crosstalks between inter-pathways/ inter-cell types by integrated machine learning algorithms.

### Case Study 1: Drug-Specific Interferon Metapathways Underlie Treatment Response Heterogeneity in RA

First, we applied Random Forests association analysis to drug response (0-nonresponse, 1-response) data after ICA in RNAcompare based on 3^rd^ level granularity defined in RNAcompare. We can see metagene_3 shows positively related to drug response in Figure 2-A. We did enrichment analysis and found it was enriched in IFN signalling in Figure 2-B. We then did 2^nd^ level ICA analysis based on metagenes which we called as metapathway analysis. Figure 2-C shows after 2^nd^ time ICA, metagenes were decomposed into metapathways for inter-pathway interactions. We can see meta_2 includes metagene_3_1st standing for metagene_3 mentioned before. Also, we think meta_2 includes related metapathways cooperating with metagene_3_1st, which are explained in Table 2-A in metapathway_2(All). Next, we used the same data and did metapathway (2^nd^ level ICA) analysis based on anti-TNF and Rituximab treatment separately, showed in Figure 2-D-E, and looked for metapathways containing metagene_3_1st. We found in Figure 2-D, metagene_3_1st belongs to meta_6, while in Figure 2-E, metagene_3_1st belongs to meta_3. We did the similar enrichment analysis for metapathway_6(Anti-TNF) and metapathway_3(Rituximab) in Table 2. Interestingly, we found the two metapathways were slightly different: metapathway for Rituximab exhibits more related to type II IFN. This may be partly because the thresholds we used to select candidate metagenes during generalised metapathway analysis. However, it cannot be fully explained by this reason because the order of the metagenes in the list also matters, since metagene_14_1st exists in Rituximab rather than Anti-TNF group. We could only contribute this to selection bias/ heterogeneity at the baseline. Figure 3-A shows SHAP importance plot for the whole dataset based on metapathways. We can see metapathway_1 is the most positively related to pain this time, while metapathway_3 exhibits complexity. When we separately plotted the importance for Rituxmab (Figure 3-B) and anti-TNF (Figure 3-C), we found metapathway_3 exhibits the characteristic of type I IFN in both plots, namely lower expression of metapathway_3 in Figure 3-B exhibits good response to Rituximab, and higher expression of metapathway_3 in Figure 3-C exhibits good response to anti-TNF, which can explain the complexity in Figure 3-A, why red dots are symmetrically distributed for metapathway_3. This seems a contradiction based on the presumption in RNAcompare that metagene_3_1st behaved more like type II IFN.

**Figure 2.**
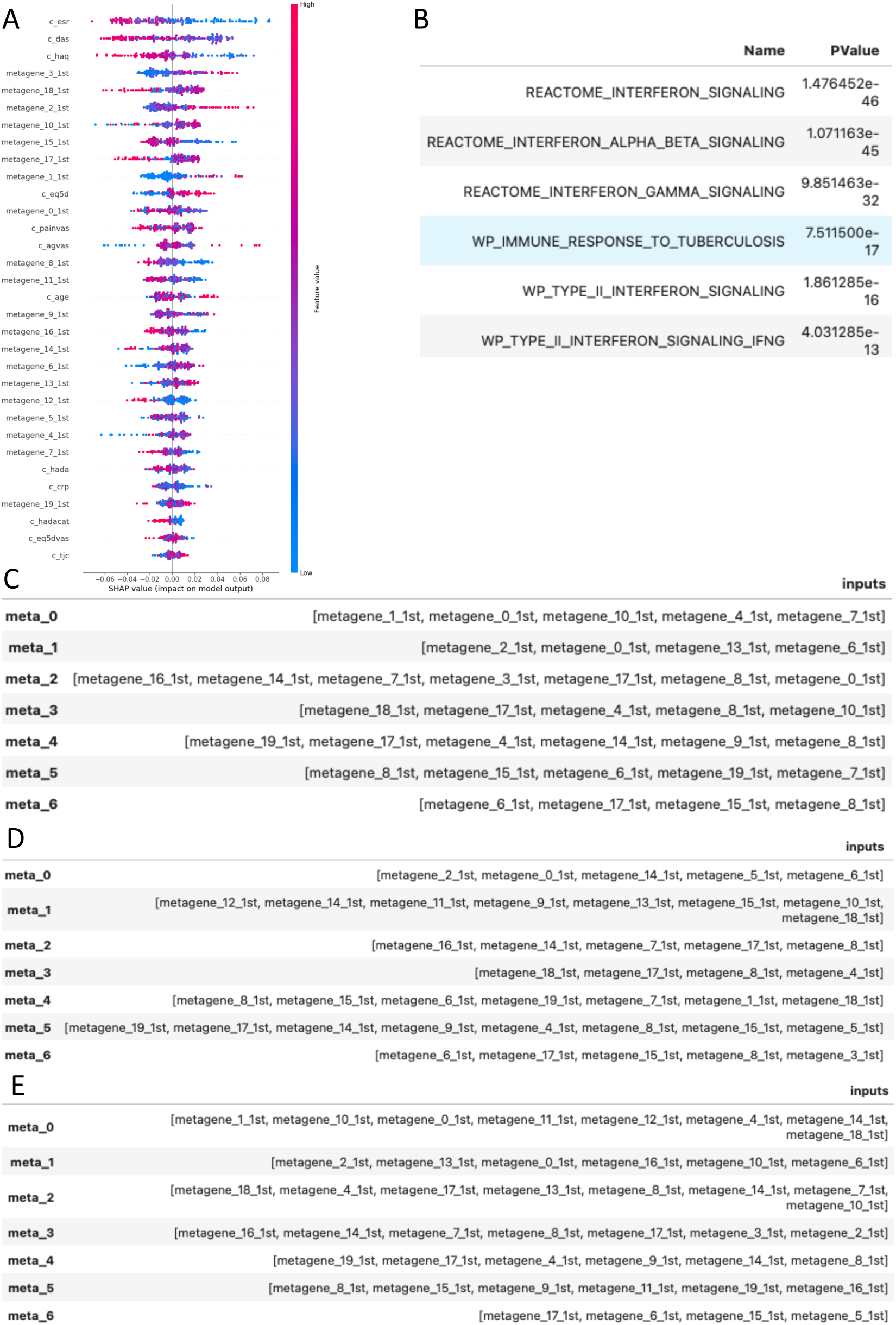
(A)SHAP importance plot for metagenes. (B) Enrichment analysis of metagene_3_1st; (C) Componets of metapathway analysis for all data; (D) Components of metapathway analysis for only Anti-TNF; (E) Components of metapathway analysis for ony Rituximab;

**Figure 3.**
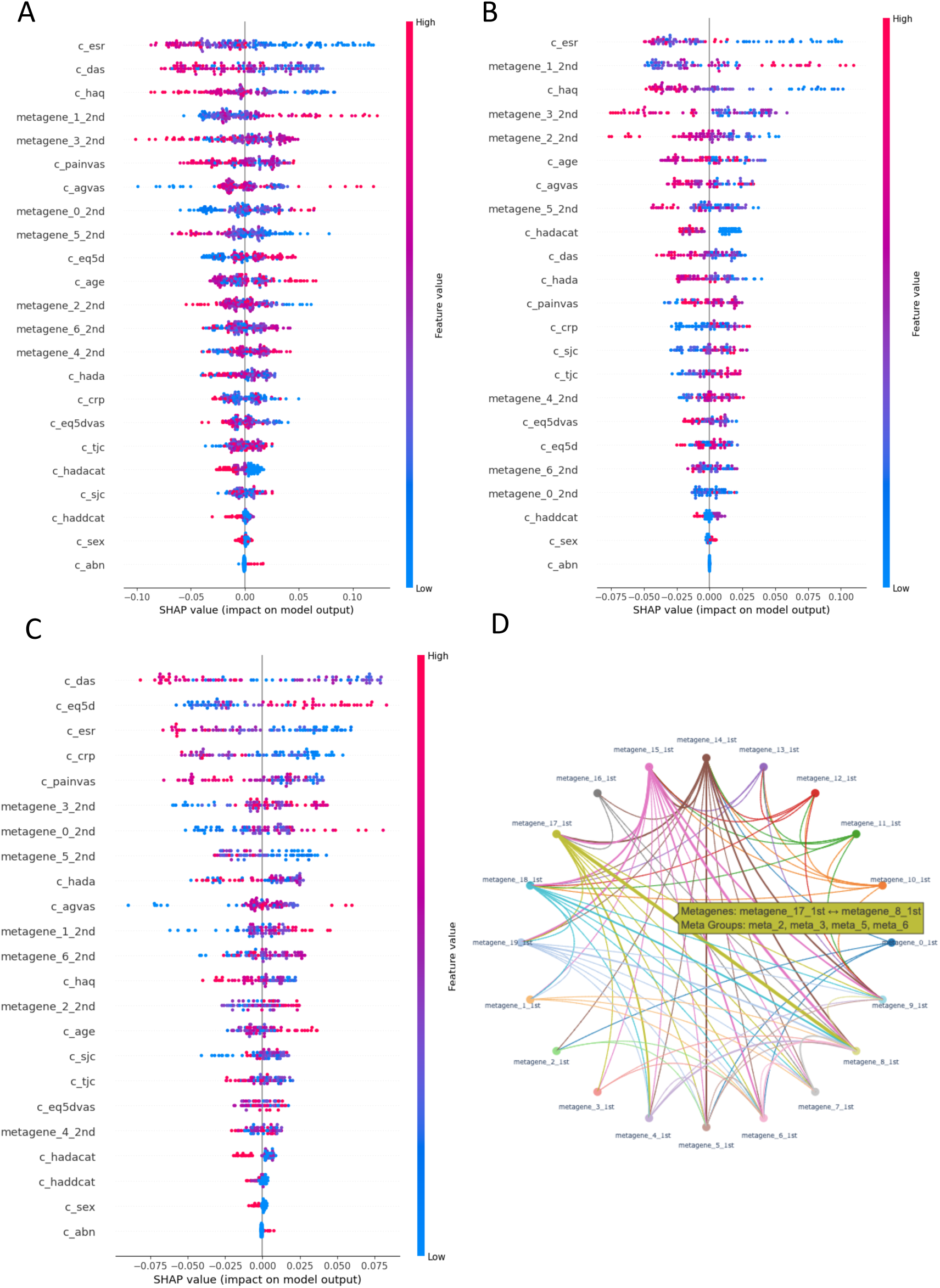
(A) SHAP importance plot for metapathways based on all data; (B) SHAP importance plot for metapathways based on Rituximab; (C) SHAP importance plot for metapathways based on anti-TNF; (D) Relationship of metagenes;

Of Note, after comparing 3 plots in Figure 3, we can see the introduction of Rituximab activates metapathway_1, making it more important to pain. Because metapathway_1 includes metagene_2_1st, which is L1CAM/Notch related pathway (see Table 2-A Metapathway_3(Rituximab))^5,35^, this explains the contradiction that during the 1^st^ ICA level analysis, the effect was contributed to metagene_3_1st for Rituximab responders.

One plausible explanation for this change is that HSP90AA1 stabilizes survival pathways in B cells, ensuring their proper function in immune regulation—notably through the Rho GTPase cycle and B cell receptor (BCR) signaling pathways^36,37^, especially in an environment with high expression of type I IFN environment. Following prior ineffective csDMARD treatments, which may have reduced type I IFN levels, the introduction of Rituximab could allow HSP90AA1 to shift its role toward stabilizing components of type II IFN signaling^36–38^, possibly amplifying inflammation and immune responses, leading to further activation of immune cells that could maintain or enhance autoimmune processes. To support this hypothesis, adding HSP90AA1 as a feature to the first-level ICA result and doing metapathway analysis again, we highlight its central regulatory role. See Appendix I.

Figure 3-D shows the relationship between metagenes based on metapathways: each circle (node) represents a metagene; each line (edge) connects metagenes within the same metapathway; the size of link is proportional to its frequency across all pathways, another perspective of visualising hub metagenes. The interaction between (metagene_17_1st, metagene_8_1st) appear frequently, representing the interaction between class I MHC mediated antigen processing presentation and B cell receptor signalling^39^; (metagene_15_1st, metagene_8_1st) represent the interaction between EJC-independent NMD and B cell signalling^40^.

## Conclusion

Through 2^nd^ level ICA analysis, we found cooperated pathways and explained the most important metapathways to the treatment outcome. Besides, we found its activated pathway after introducing Rituximab.

### Case Study 2: Immune Feedback Loops to pain at transcriptomic level: The Role of Type II IFN and Complement Cascade

In this case, we will adopt Causal Forests based on cohort-ICA similar to cases in RNAcompare for the combination of ORBIT and PEAC. But this time we didn’t use Pearson Correlation matrix for variable selection.

Figure 4-A shows the SHAP importance of metapathways to pain. Figure 4-B shows their components after Enrichment analysis. (metagene_0_orb, metagene_3_pea) grouped together in metapathway_2 as expected because they are identical.

**Figure 4.**
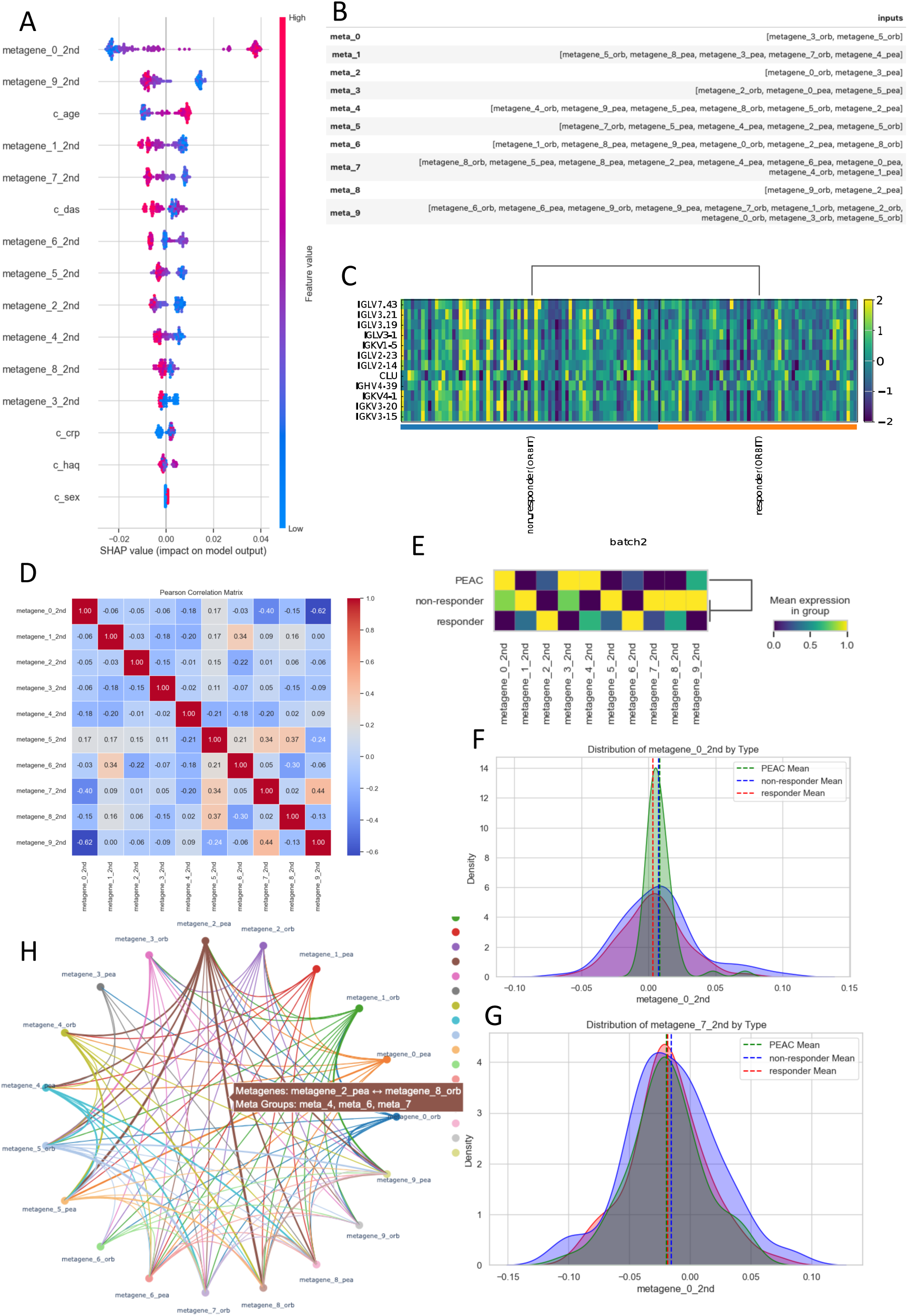
(A) SHAP importance of metapathways; (B) Componets of metapathways; (C) Signatures of complement cascade pathway; (D) Pearson Correlation Matrix of metapathways; (E) Matrixplot of 3 batches scaled by metagenes; (F) Distribution of metapathway_0 of 3 batches; (G) Distribution of metapathway_7 of 3 batches. (H) Relationship of metagenes.

We can see the most important metapathway is metapathway_0, including 2 metagenes, metagene_3_orb and metagene_5_orb. The 2^nd^ important metapathway is metapathway_9. After that we noticed complement cascade in metapathway_1 acting as a feedback hub, and linking innate immune responses (alpha defensins) and coagulation/thrombosis (platelet activation).

C3 and C5 activation are crucial steps in the complement cascade, and their activation has significant implications for immune responses, particularly in inflammation and immune complex-mediated diseases like RA^41^. Using RNAcare, we drew the heatmap plot for ORBIT. As we expected in Figure 4-C, non-respondent group shows high expression with signatures of complement cascade pathway, identical with the result in RNAcompare.

Next, we drew a Pearson Correlation Matrix to explore the relationship among metapathways in Figure 4-D. We can see metapathway_0 and metapathway_9 are negatively correlated. Notably, metapathway_9 is enriched in type II IFN. Therefore, our assumption is under Rituximab and its induction of type II IFN, metapathway_9 proliferated the damage from metapathway_0, while relieving the pain.

To further prove this assumption, we did a matrixplot scaled by each pathway in Figure 4-E. As we see, the mean expression of metapathway_0 in responder group (ORBIT) is lower than that in non-responder group (ORBIT). Since we already known that responder group had higher expression of type II IFN (RNAcompare), the result proves our assumption. Besides, we also see metapathway_7 (complement cascade related pathway) is higher in non-response group. We can also prove it in density plots in Figure 4-FG.

Different from non-responders, for Rituximab responders in metapathway_6, the participation of type II IFN can activate the cooperation of IL10 signaling and complement cascade in the negative feedback loop, relieving the pain from metapathway_7^42,43^.

Figure 4-E shows the interactions between metagenes. (metagene_2_pea, metagene_8_orb) represents the most frequent one between antigen presentation and TLR signaling^44^; (metagene_2_pea, metagene_5_pea) stands for the one between nkcells pathway and alpha defensin^45^.

Besides, we also think the involvement of type II IFN have two roles: on one hand, they can drive pain progression within the same level, to make the tissue less damaged; on the other hand, they can act in a negative feedback loop to prevent excessive inflammation and pain sensitization between different layers (tissues/omics layers).

### Case Study 3: Synovium vs. Blood: Distinct Immune Feedback Loops in RA Pain Regulation

In this case, we applied Causal Forests, connecting different tissues to pain using PEAC. Figure 5-A shows SHAP importance. Figure 5-B shows metagene members for each metapathways. Figure 5-C shows the relationships among metapathways. We can see Group 1: (metapathway_3, metapathway_4, metapathway_5, metapathway_8, metapathway_9) negatively correlated with Group 2: (metapathway_1, metapathway_2). Figure 5-D shows the matrix plot scaled by tissue types.

**Figure 5.**
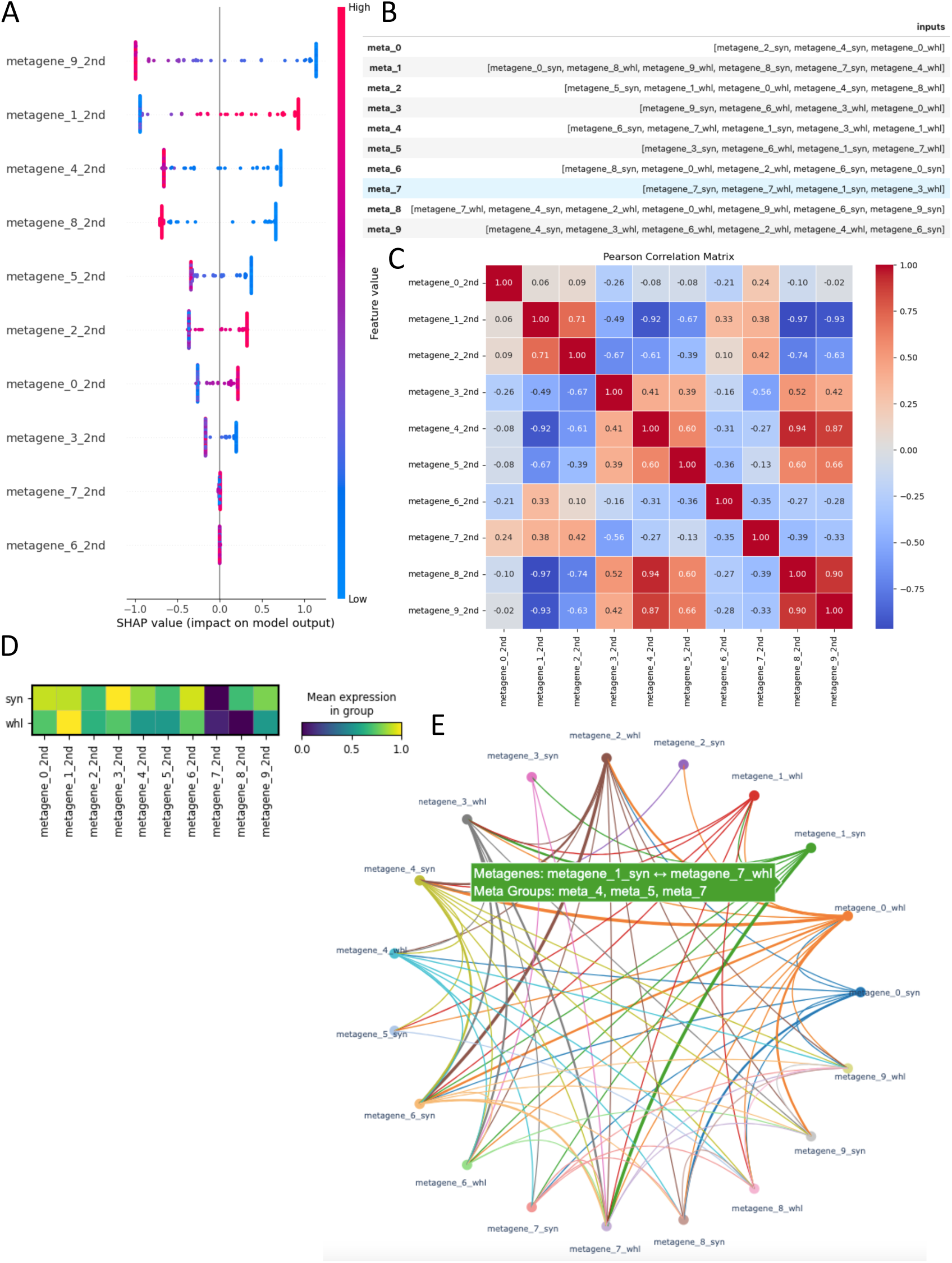
(A) SHAP importance plot; (B) Components of metapathway analysis; (C) Pearson Correlation Matrix of metapathways; (D) Matrix plot scaled by batches; (E) Relationship f metagenes.

Combining Figure 5-CD, we can see Group1 is more active at synovium level while Group 2 is more active at whole blood level.

- **Group 1 (G1)** - Synovium-level G1 consists of pathways related to growth factor signaling, PI3K activation, synovial remodeling, immune activation, and nerve interaction, all of which can contribute to pain in RA:

1. From metapathway_4 and metapathway_5, we know that hub metapathways are more related to PI3K signalling, antimicrobial peptides, and fibrin clot formation.
2. metapathway_4 and metapathway_5 are the 1^st^ level proliferation including IL-1 pathway; metapathway_3 is 2^nd^ level proliferation including type II IFN; metapathway_9 and metapathway_8 are the positive feedback loops;
3. metapathway_3 is the most enriched in synovium tissue in Figure 5-D, including nervous system development, which means the proliferation from synovium to the nerve.
- **Group 2 (G2)** - Blood-level

1. metapathway_1 is the proliferation and metapathway_2 is the negative feedback loop;
2. REACTOME_TNFR2_NON_CANONICAL_NF_KB_PATHWAY Non-canonical NF-κB signaling via TNFR2 is often protective like IL-10, especially in Tregs (Regulatory T Cells) and immune resolution mechanisms to supports Tregs’ suppressive function, while activating PI3K/Akt signaling^46^.
3. WP_INFLAMMATORY_RESPONSE_PATHWAY This pathway may buffer excessive inflammation by regulating cytokine signaling. Instead of promoting chronic inflammation, it helps restore immune balance, potentially reducing pain.
4. WP_QUERCETIN_AND_NFKB_AP1_INDUCED_APOPTOSIS Pro-apoptotic mechanisms in blood may limit inflammatory cell survival in synovium, reducing pain-driving immune responses.
5. PID_INTEGRIN4_PATHWAY & WP_BURN_WOUND_HEALING Involved in tissue repair and immune regulation, potentially counteracting chronic inflammation in the synovium^47^.

G2 appears to have immune-regulatory, apoptotic, and anti-inflammatory properties that could act as a compensatory mechanism to suppress excessive inflammation and pain from G1. G1 pathways in the synovium drive inflammation and pain, while G2 pathways in blood counteract excessive inflammation and promote immune regulation. This suggests that therapies enhancing G2 (e.g., TNFR2 agonists, apoptosis inducers, or regulatory immune modulators) could help reduce RA pain. The inflammatory environment in the synovium allows for positive feedback loops, where complement activation enhances PI3K signaling, promoting fibroblast proliferation and tissue inflammation. This feedback contributes to the chronic inflammation. In the bloodstream, the immune system is more regulated by mechanisms like Tregs. Here, TNFR2 signaling enhances Treg expansion, which suppresses complement activation and PI3K signaling to prevent excessive immune responses, thus creating a negative feedback loop. This difference in the feedback loops can be understood as a tissue-specific response to the immune system’s need for either promotion of inflammation (in synovium) or immune regulation (in blood). Until now, we can explain the phenomenon in RNAcare why L1CAM is downregulated to pain in blood, but upregulated to pain in DRG.

Besides, we found in terms of crosstalk, CCL2 recruits monocytes/macrophages from the bloodstream into inflamed tissues, including the synovium in RA, acting as a bridge from the bloodstream to the synovium, drawing immune cells into the joint space, thereby initiating and amplifying inflammation; CCL5 promotes the retention and activation of T cells and macrophages in the synovium, sustaining inflammation, acting as a feedback signal, maintaining chronic inflammation by keeping immune cells in the synovium and further activating them^48^. Hence, Targeting CCL2 (blocking monocyte recruitment) might help reduce immune cell infiltration; Targeting CCL5 (disrupting retention and activation of immune cells) could break the chronic inflammation loop. And during the feedback loop CCL5 may potentially increase the risk of getting pancreatic cancer for RA patients^49^.

Meanwhile, we found the progression of pain in RA following the sequence: blood → synovium → dorsal root ganglion (DRG). The CCL2/CCR2 axis has been implicated in peripheral inflammatory pain sensitization. Research indicates that CCL2 can act directly on DRG neurons, leading to increased neuronal excitability and pain hypersensitivity^50^.

Finally, Figure 5-E shows the interaction plot among metagenes based on Figure 5-B. As we see, (metagene_1_syn, metagene_7_whl) crosstalks frequently, representing the crosstalk between antimicrobial peptides (alpha defensins) in blood and PI3K/AKT signalling in synovium^51^; (metagene_2_whl, metagene_6_syn) crosstalks frequently, representing the crosstalk between glucocorticoid receptor pathway in blood and O-linked glycosylation in synovium^52^; (metagene_0_whl, metagene_4_syn) crosstalks frequently, representing the crosstalk between IFN signaling and complement system pathway/ medicus reference GF RTK PI3K signalling^53^.

## Conclusion

Through combining different tissues, we know the relationship between different metapathways, the progression direction related to pain, the mechanism for RA to recruit/retain immune cells and the hub metapathways.

### Case Study 4: Cross-Omics Regulation of Pain: The Role of type II IFN, IL-6, and IL-10 in RA

In this case, we applied Causal Forests, connecting different omics data to pain using RA-MAP.

Figure 6-A shows SHAP importance. Figure 6-B shows metagene members for each metapathways. Figure 6-C shows the relationships among metapathways. We can see G1: (metapathway_0, metapathway_9); G2: (metapathway_6, metapathway_7, metapathway_8); G3: (metapathway_1, metapathway_2, metapathway_3, metapathway_4). Figure 6-D shows the matrix plot scaled by omics types.

**Figure 6.**
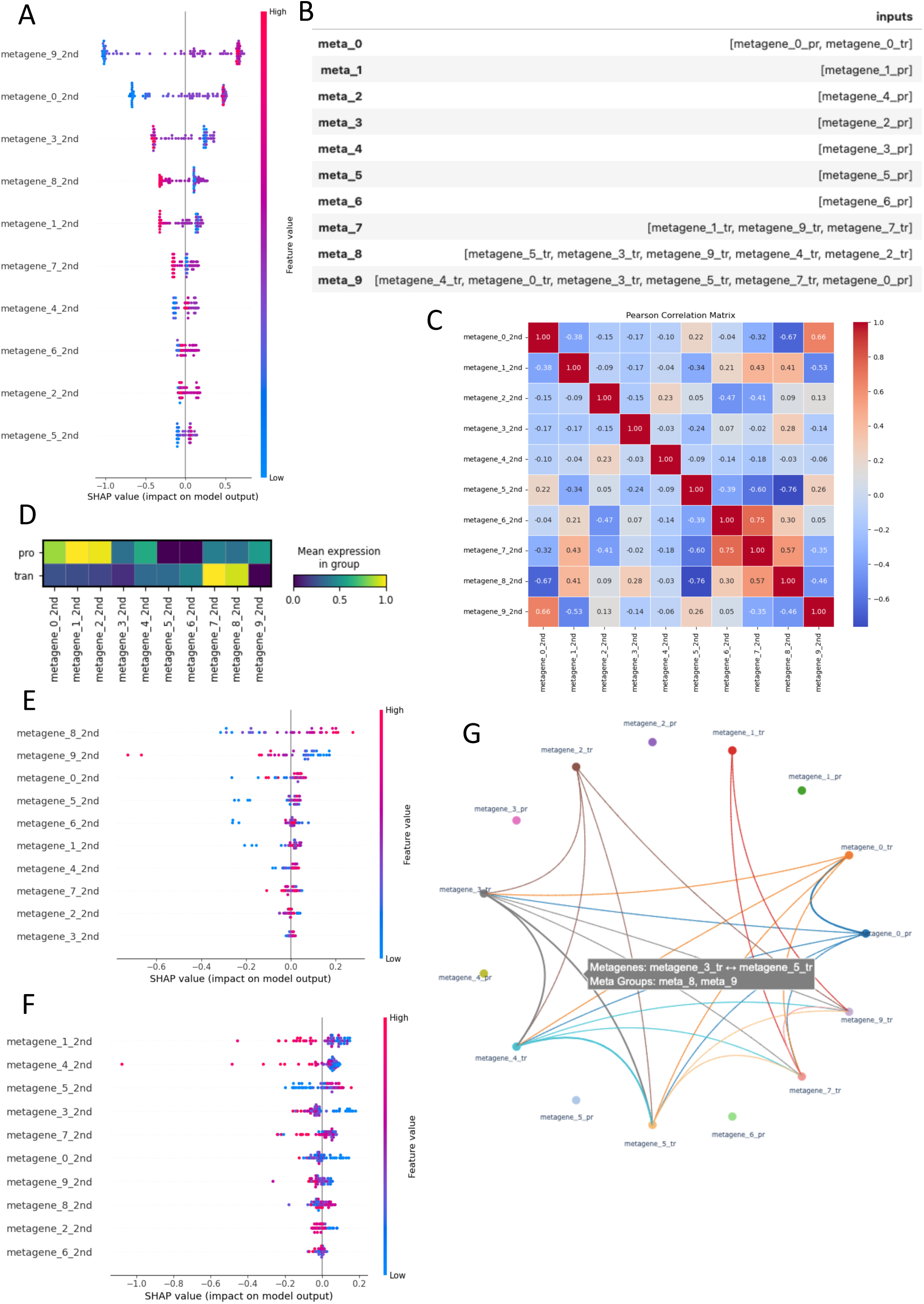
(A) SHAP importance plot; (B) Components of metapathway analysis; (C) Pearson Correlation Matrix of metapathways; (D) Matrix plot scaled by batches; (E) SHAP importance plot for only transcriptomic level based on association analysis; (F) SHAP importance plot for only proteomic level based on association analysis; (G) Relationship of metagenes.

Combining Figure 6-CD, we can see in proteomics is active in G3 and transcripts is active in G2. Given the relevance to pain, we propose the following explanation:

#### metapathway_0 as a Hub Metapathway

It contains platelet activation, fibrin clot formation, and peptide ligand binding receptor pathways, which are highly interconnected with inflammatory and immune responses. Platelet activation and coagulation (e.g., GPVI-mediated activation cascade, intrinsic pathway of fibrin clot formation) are key contributors to inflammation and pain in RA.

This suggests it acts as a central regulatory network for inflammatory and immune activation, which then influences metapathway_9, metapathway_8, and metapathway_1.

#### metapathway_9 (Inflammation and Immune Activation)

Promoted by metapathway_0, antimicrobial peptides and NF-κB activation (via TRAF6) indicate a strong link to innate immune responses and chronic inflammation, which drive pain.

#### metapathway_8 (Adaptive Immunity and Chronic Inflammation)

TNFR2 and T-cell receptor signaling point to adaptive immune activation, contributing to sustained inflammation and blood damage in RA; NF-κB activation (TRAF6-mediated and non-canonical TNFR2)^54^ and type II IFN^55^ links it back to pain sensitization.

#### metapathway_1 (Resolution Mechanism via IL-6 and Fibrin Clot Formation)

IL-6 signaling and fibrin clot formation suggest a role in transitioning inflammation toward resolution, aligning with the crosstalk between IL-6 and IFN-γ in immune regulation^56,57^.

#### metapathway_7 (Adaptive Immunity and Chronic Inflammation)

It shares similarities with metapathway_8.

#### (metapathway_8 & metapathway_7) & metapathway_1 act as complementary modulators to metapathway_0 & metapathway_9 for transcriptomic and proteomic level respectively

type II IFN (transcriptomic level) and IL-6 (proteomic level) regulate the immune response, acting as negative feedback loop and preventing excessive pain sensitization while TNFR2 non-canonical NF-κB pathway supports Tregs’ suppressive function; Therefore, metapathway_1 is the pain buffer at proteomic level and metapathway_7& metapathway_8 is the pain buffer at transcriptomic level.

#### Metagene_0_pr was the only hub metagene at proteomics level communicating with transcriptomic level

Figure 6-E is the SHAP importance for only transcriptomic level and Figure 6-F is the SHAP importance for only proteomic level. As what we expected before, in Figure 6-E, metapathway_8 buffered metapathway_9 and metapathway_0, contributing the most pain; while in Figure 6-F, metapathway_1and metapathway_4 buffered metapathway_9 and metapathway_0. The negative feedback loop at different omics levels is a protective mechanism.

Figure 6-G shows the interaction plot among metagenes based on Figure 6-B: (metagene_3_tr, metagene_5_tr) represents interaction between TRAF6 mediated NFKB activation and cytokine receptor signaling^58^.

Interestingly, we notice the different roles between metapathway_1 and metapathway_4 in the two omics lays in Figure 6-EF to pain. It is because they act as pain regulators not only at the same omics level but also across levels. Then, we checked the enrichment result of metapathway_4 which is enriched in IL-10 signaling. IL-10 signaling is a well-known negative feedback mechanism that dampens excessive inflammation, which could prevent sustained pain sensitization^59^.

## Conclusion

Through combining different omics data, we know the relationship between different metapathways correlated to pain. At the transcriptomic level, type II IFN often acts as a negative feedback loop by inducing immune-modulatory responses; at the proteomic level, IL-6 and IL-10 appear to play a more dominant role in negative feedback compared to type II IFN. We can see in blood tissue, feedback loop is always negative across different omics. The pain buffer serves as a protective mechanism to modulate pain, helping to maintain balance and prevent excessive pain perception. If this buffer is removed or disrupted, the body can lose its ability to regulate pain properly, leading to increased pain sensitivity or chronic pain. Many pain treatments aim to target this buffer but must do so carefully to avoid removing protective feedback because pain in this situation is helpful to prevent further tissue damage.

### Case Study 5: Divergent Buffering Strategies in RA and HF: The Role of IFN Signaling in Disease Severity

In this case, we applied Causal Forests, connecting different diseases to disease score using HF and RA data.

Figure 7-A shows SHAP importance. Figure 7-B shows metagene members for each metapathways. Figure 7-C shows the relationships among metapathways. We can see G1: (metapathway_0, metapathway_9), G2: (metapathway_1, metapathway_8) are negatively correlated. Figure 7-D shows the matrix plot scaled by disease types. Combing Figure 7-A and Table 6, we know the hub pathways is (metapathway_6, metapathway_7) (HF related), (alpha defensins, REACTOME_RESPONSE_TO_ELEVATED_PLATELET_CYTOSOLIC_CA2) (RA related) and metapathway_0 (shared). Although metapathway_6 and metapathway_7 shared almost the same metapathways, ICA separates them as different components, which will be analysed later; metapathway_9 further proliferates metapathway_0 in both RA and HF by IFN signaling; For RA only, metapathway_1 and metapathway_8 are negatively related to metapathway_0 because metapathway_9 suppressed their interactions with metapathway_0 (Table 3) and they can only be proliferated by type II IFN (6_op, 7_op).

**Figure 7.**
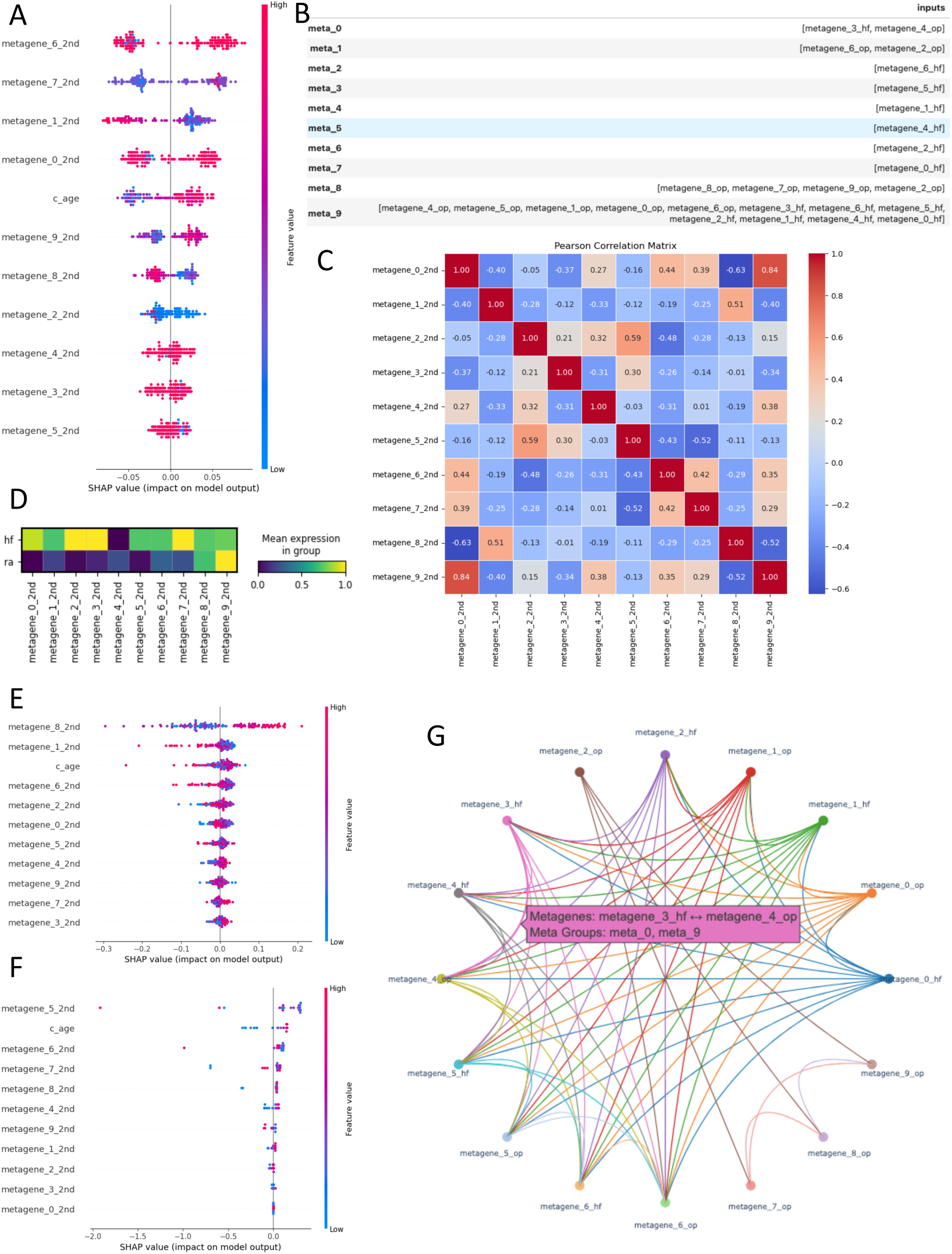
(A) SHAP importance plot; (B) Components of metapathway analysis; (C) Pearson Correlation Matrix of metapathways; (D) Matrix plot scaled by diseases; (E) SHAP importance plot for only RA based on association analysis; (F) SHAP importance plot for only HF based on association analysis; (G) Relationship of metagenes.

**Table 3.**
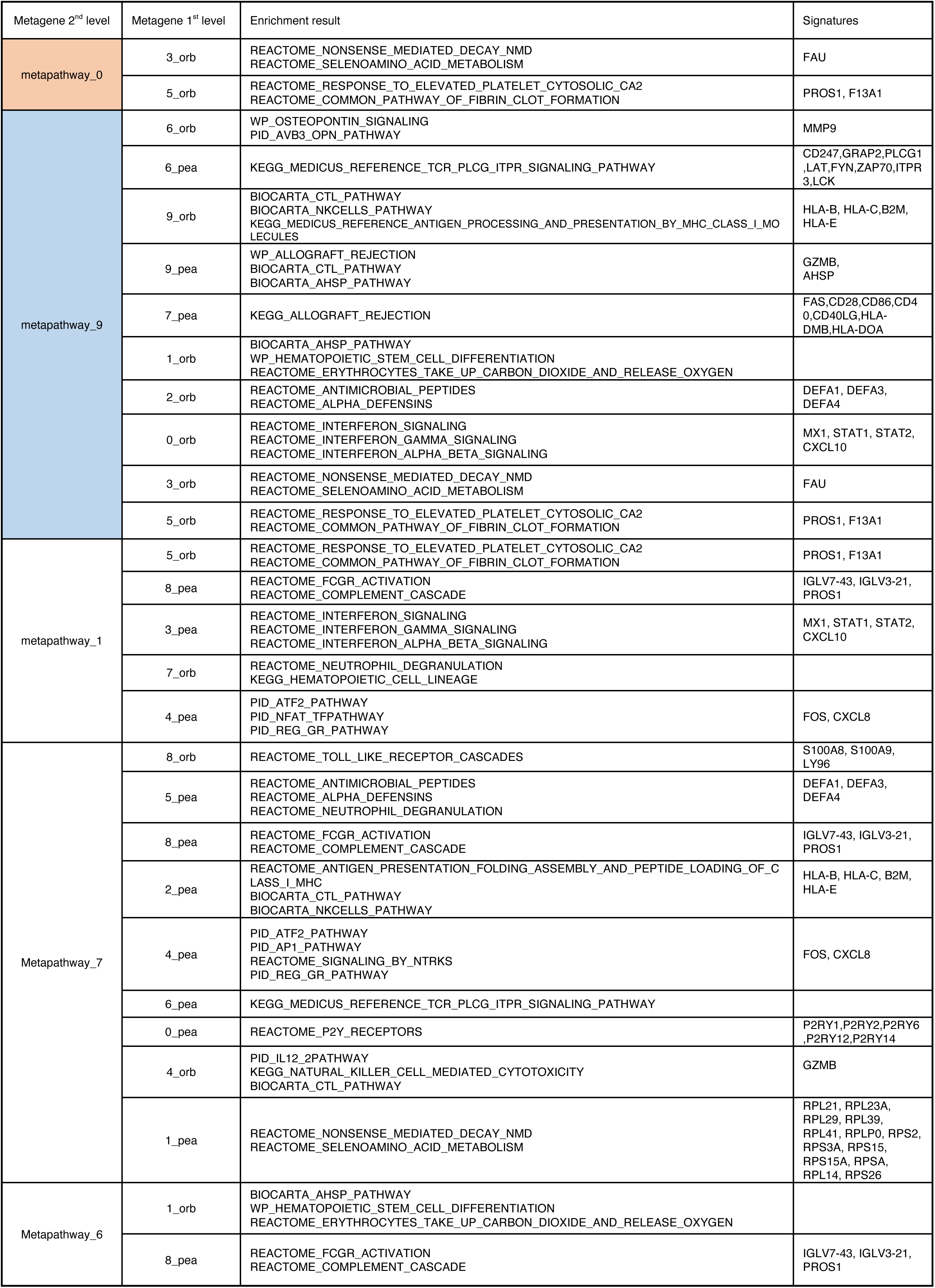
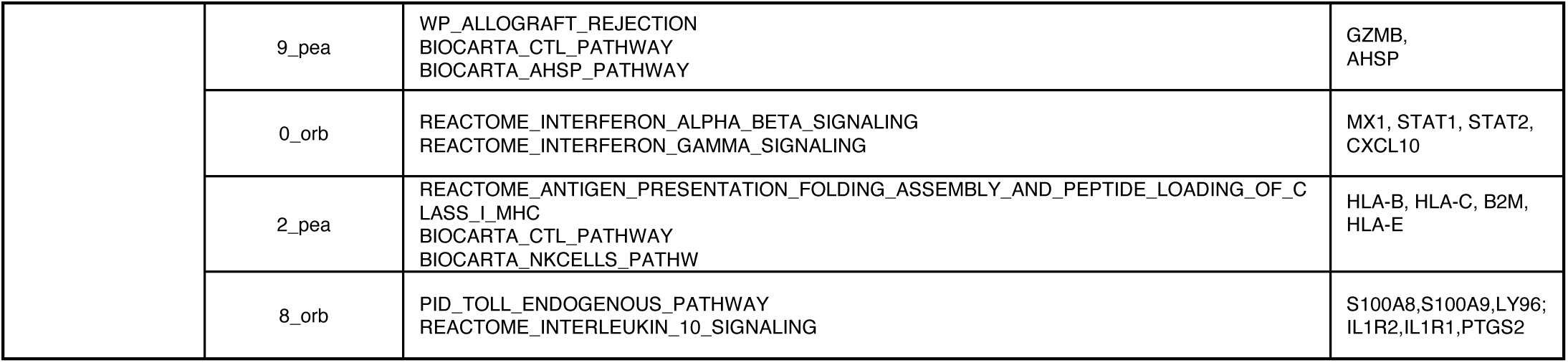
Metapathway analysis.

**Table 4.** Metapathway analysis.

| metagene_2nd_level | metagene_1st_level | Enrichment result | Signatures |
| --- | --- | --- | --- |
| metapathway_9 | 4_syn | KEGG_MEDICUS_REFERENCE_GF_RTK_PI3K_SIGNALING_PATHWAY<br>WP_COMPLEMENT_SYSTEM<br>WP_COMPLEMENT_AND_COAGULATION_CASCADES | EREG<br>CFI,CFD,C4A,ADIPOQ,<br>CFI,F3,CFD |
|  | 3_whl | REACTOME_ANTIMICROBIAL_PEPTIDES | CTSG, LTF |
|  | 6_whl | WP_NEURAL_CREST_CELL_MIGRATION_DURING_DEVELOPMENT<br>REACTOME_NERVOUS_SYSTEM_DEVELOPMENT | EPHB4,MMP9 |
|  | 2_whl | BIOCARTA_CCR5_PATHWAY<br>WP_BURN_WOUND_HEALING | CCL2 |
|  | 4_whl | REACTOME_NEUTROPHIL_DEGRANULATION<br>KEGG_MEDICUS_REFERENCE_ANTIGEN_PROCESSING_AND_PRESENTATION_BY_MHC_CLASS_I_MOLECULES<br>BIOCARTA_CTL_PATHWAY | B2M |
|  | 6_syn | REACTOME_O_LINKED_GLYCOSYLATION<br>PID_INTEGRIN1_PATHWAY<br>REACTOME_RESPONSE_TO_ELEVATED_PLATELET_CYTOSOLIC_CA2 | THBS1 |
| metapathway_1 | 0_syn | REACTOME_TNFR2_NON_CANONICAL_NF_KB_PATHWAY<br>WP_INFLAMMATORY_RESPONSE_PATHWAY<br>KEGG_CYTOKINE_CYTOKINE_RECEPTOR_INTERACTION | TNFRSF13C, BIRC3,<br>CD27, TNFRSF18,<br>ZAP70, LCK,<br>CCL5 |
|  | 8_whl | BIOCARTA_AHSP_PATHWAY<br>REACTOME_ERYTHROCYTES_TAKE_UP CARBON_DIOXIDE_AND_RELEASE_OXYGEN<br>WP_QUERCETIN_AND_NFKB_AP1_INDUCED_APOPTOSIS | HBA1, PTGS2 |
|  | 9_whl | WP_INTERACTIONS_OF_NATURAL_KILLER_CELLS_IN_PANCREATIC_CANCER | KLRK1, CCL5 |
|  | 8_syn | WP_TOLLLIKE_RECEPTOR_SIGNALING<br>WP_OSTEOCLAST_SIGNALING | CXCL10,SPP1 |
|  | 7_syn | PID_INTEGRIN4_PATHWAY<br>WP_BURN_WOUND_HEALING | LAMA1 |
|  | 4_whl | KEGG_MEDICUS_REFERENCE_ANTIGEN_PROCESSING_AND_PRESENTATION_BY_MHC_CLASS_I_MOLECULES<br>BIOCARTA_CTL_PATHWAY | B2M |
| metapathway_4 | 6_syn | REACTOME_O_LINKED_GLYCOSYLATION<br>PID_INTEGRIN1_PATHWAY<br>REACTOME_RESPONSE_TO_ELEVATED_PLATELET_CYTOSOLIC_CA2 | THBS1 |
|  | 7_whl | REACTOME_NEGATIVE_REGULATION_OF_THE_PI3K_AKT_NETWORK<br>WP_PI3KAKT_SIGNALING | FGF23,NRG1,FLT3 |
|  | 1_syn | REACTOME_ANTIMICROBIAL_PEPTIDES<br>PID_IL1_PATHWAY<br>WP_PHOTODYNAMIC_THERAPYINDUCED_NFKB_SURVIVAL_SIGNALING | S100A9,PI3,LCN2,<br>IL1B |
|  | 3_whl | REACTOME_ANTIMICROBIAL_PEPTIDES | CTSG,LTF,LCN2 |
|  | 1_whl | REACTOME_RESPONSE_TO_ELEVATED_PLATELET_CYTOSOLIC_CA2<br>REACTOME_FORMATION_OF_FIBRIN_CLOT_CLOTTING_CASCADE | PROS1 |
| metapathway_8 | 7_whl | REACTOME_NEGATIVE_REGULATION_OF_THE_PI3K_AKT_NETWORK<br>WP_PI3KAKT_SIGNALING | FGF23,NRG1,FLT3 |
|  | 4_syn | KEGG_MEDICUS_REFERENCE_GF_RTK_PI3K_SIGNALING_PATHWAY<br>WP_COMPLEMENT_SYSTEM<br>WP_COMPLEMENT_AND_COAGULATION_CASCADES | EREG<br>CFI,CFD,C4A,<br>ADIPOQ,<br>CFI,F3,CFD |
|  | 2_whl | WP_BURN_WOUND_HEALING<br>PID_TCR_CALCIUM_PATHWAY<br>WP_GLUCCORTICOID_RECEPTOR_PATHWAY | CCL2, JUN,PTGS2 |
|  | 0_whl | WP_TYPE_II_INTERFERON_SIGNALING<br>REACTOME_INTERFERON_ALPHA_BETA_SIGNALING<br>WP_SPINAL_CORD_INJURY | STAT1, ISG15,<br>OAS1, CXCL10,<br>CCL2, TNFSF13B,<br>CXCL10 |
|  | 9_whl | WP_INTERACTIONS_OF_NATURAL_KILLER_CELLS_IN_PANCREATIC_CANCER | KLRK1,CCL5 |
|  | 6_syn | REACTOME_O_LINKED_GLYCOSYLATION<br>PID_INTEGRIN1_PATHWAY<br>REACTOME_RESPONSE_TO_ELEVATED_PLATELET_CYTOSOLIC_CA2 | THBS1 |
|  | 9_syn | REACTOME_INITIAL_TRIGGERING_OF_COMPLEMENT<br>KEGG_COMPLEMENT_AND_COAGULATION_CASCADES<br>WP_T_CELL_MODULATION_IN_PANCREATIC_CANCER | C1QA,CFD,TNFRSF4,LGALS9<br>KLRK1,CCL5 |
| metapathway_5 | 3_syn | REACTOME_ERBB2_ACTIVATES_PTK6_SIGNALING<br>REACTOME_CONSTITUTIVE_SIGNALING_BY_ABERRANT_PI3K_IN_CANCER | ERBB3,EREG,PTK6 |
|  | 6_whl | WP_NEURAL_CREST_CELL_MIGRATION_DURING_DEVELOPMENT<br>REACTOME_NERVOUS_SYSTEM_DEVELOPMENT | EPHB4,MMP9 |
|  | 1_syn | REACTOME_ANTIMICROBIAL_PEPTIDES<br>PID_IL1_PATHWAY<br>WP_PHOTODYNAMIC_THERAPYINDUCED_NFKB_SURVIVAL_SIGNALING | S100A9,PI3,LCN2,IL1B |
|  | 7_whl | REACTOME_NEGATIVE_REGULATION_OF_THE_PI3K_AKT_NETWORK<br>WP_PI3KAKT_SIGNALING | FGF23,NRG1,FLT3 |
| metapathway_2 | 5_syn | REACTOME_ACTIVATED_NOTCH1_TRANSMITS_SIGNAL_TO_THE_NUCLEUS<br>REACTOME_PLATELET_ADHESION_TO_EXPOSED_COLLAGEN | JAG2,NOTCH1<br>VWF,GP1BA |
|  | 1_whl | REACTOME_RESPONSE_TO_ELEVATED_PLATELET_CYTOSOLIC_CA2<br>REACTOME_FORMATION_OF_FIBRIN_CLOT_CLOTTING_CASCADE | PROS1 |
|  | 0_whl | WP_TYPE_II_INTERFERON_SIGNALING<br>REACTOME_INTERFERON_ALPHA_BETA_SIGNALING<br>WP_SPINAL_CORD_INJURY | STAT1, ISG15,<br>OAS1, CXCL10,<br>CCL2,<br>TNFSF13B,CXCL10 |
|  | 4_syn | KEGG_MEDICUS_REFERENCE_GF_RTK_PI3K_SIGNALING_PATHWAY<br>WP_COMPLEMENT_SYSTEM<br>WP_COMPLEMENT_AND_COAGULATION_CASCADES | EREG<br>CFI,CFD,C4A,ADIPOQ,<br>CFI,F3,CFD |
|  | 8_whl | BIOCARTA_AHSP_PATHWAY<br>REACTOME_ERYTHROCYTES_TAKE_UP CARBON_DIOXIDE_AND_RELEASE_OXYGEN<br>WP_QUERCETIN_AND_NFKB_AP1_INDUCED_APOPTOSIS | HBA1, PTGS2 |
| Metapathway_3 | 9_syn | REACTOME_INITIAL_TRIGGERING_OF_COMPLEMENT<br>KEGG_COMPLEMENT_AND_COAGULATION_CASCADES<br>WP_T_CELL_MODULATION_IN_PANCREATIC_CANCER | C1QA,CFD,TNFRSF4,LGALS9<br>KLRK1,CCL5 |
|  | 6_whl | WP_NEURAL_CREST_CELL_MIGRATION_DURING_DEVELOPMENT<br>REACTOME_NERVOUS_SYSTEM_DEVELOPMENT | EPHB4,MMP9 |
|  | 3_whl | REACTOME_ANTIMICROBIAL_PEPTIDES | CTSG, LTF |
|  | 0_whl | WP_TYPE_II_INTERFERON_SIGNALING<br>REACTOME_INTERFERON_ALPHA_BETA_SIGNALING<br>WP_SPINAL_CORD_INJURY | STAT1, ISG15,<br>OAS1, CXCL10,<br>CCL2, TNFSF13B,<br>CXCL10 |

**Table 5.** Metapathway analysis.

| metagene_2<br>nd_level | metagene_1<br>st_level | Enrichment result | Signatures |
| --- | --- | --- | --- |
| metapathway_9 | 4_tr | REACTOME_ANTIMICROBIAL_PEPTIDES | CTSG,ELANE,LTF,PRTN3,LCN2,BPI |
|  | 0_tr | REACTOME_RESPONSE_TO_ELEVATED_PLATELET_CYTOSOLIC_CA2<br>REACTOME_INTRINSIC_PATHWAY_OF_FIBRIN_CLOT_FORMATION | SELP,PF4,THBS1,SPARC,VWF,ITGA2B,PROS1,SERPINE2,VWF,GP1BA |
|  | 3_tr | REACTOME_REGULATION_OF_SIGNALING_BY_CBL<br>REACTOME_TRAF6_MEDIATED_NF_KB_ACTIVATION | YES1,PIK3CA, HMGB1,S100A12 |
|  | 5_tr | KEGG_CYTOKINE_CYTOKINE_RECEPTOR_INTERACTION | CCL23,LEP,IL10,TNFSF15 |
|  | 7_tr | BIOCARTA_ARF_PATHWAY<br>REACTOME_GPVI_MEDIATED_ACTIVATION_CASCADE | PIK3CA,PIK3CG |
|  | 0_pr | REACTOME_RESPONSE_TO_ELEVATED_PLATELET_CYTOSOLIC_CA2<br>REACTOME_INTRINSIC_PATHWAY_OF_FIBRIN_CLOT_FORMATION<br>REACTOME_PEPTIDE_LIGAND_BINDING_RECEPTORS | PPBP,PDGFA,PDGFB,SPARC,TF,PF4,APP,CCL5 |
| metapathway_0 | 0_pr | REACTOME_RESPONSE_TO_ELEVATED_PLATELET_CYTOSOLIC_CA2<br>REACTOME_INTRINSIC_PATHWAY_OF_FIBRIN_CLOT_FORMATION<br>REACTOME_PEPTIDE_LIGAND_BINDING_RECEPTORS | PPBP,PDGFA,PDGFB,SPARC,TF,PF4,APP,CCL5 |
|  | 0_tr | REACTOME_RESPONSE_TO_ELEVATED_PLATELET_CYTOSOLIC_CA2<br>REACTOME_INTRINSIC_PATHWAY_OF_FIBRIN_CLOT_FORMATION | SELP,PF4,THBS1,SPARC,VWF,ITGA2B,PROS1,SERPINE2,VWF,GP1BA |
| metapathway_3 | 2_pr | WP_EXTRAFOLLCULAR_AND_FOLLICULAR_B_CELL_ACTIVATION_BY_SARSCOV2 | BCL6,SLAMF7 |
| metapathway_8 | 5_tr | KEGG_CYTOKINE_CYTOKINE_RECEPTOR_INTERACTION | CCL23,LEP,IL10,TNFSF15 |
|  | 3_tr | REACTOME_REGULATION_OF_SIGNALING_BY_CBL<br>REACTOME_TRAF6_MEDIATED_NF_KB_ACTIVATION | YES1,PIK3CA, HMGB1,S100A12 |
|  | 9_tr | REACTOME_TNFR2_NON_CANONICAL_NF_KB_PATHWAY<br>KEGG_T_CELL_RECEPTOR_SIGNALING_PATHWAY | CD27,CD40LG,PLCG1 |
|  | 4_tr | REACTOME_ANTIMICROBIAL_PEPTIDES | CTSG,ELANE,LTF,PRTN3,LCN2,BPI |
|  | 2_tr | WP_ALLOGRAFT_REJECTION | IFNG,FASLG,GNLY,GZMB |
| metapathway_1 | 1_pr | REACTOME_COMMON_PATHWAY_OF_FIBRIN_CLOT_FORMATION<br>REACTOME_FORMATION_OF_FIBRIN_CLOT_CLOTTING_CASCADE<br>WP_IL6_SIGNALING | SERPINA5,FGA,MAP2K4,JAK2 |
| metapathway_7 | 1_tr | REACTOME_ACTIVATED_TAK1_MEDIATES_P38_MAPK_ACTIVATION | UBB,MAP2K3 |
|  | 9_tr | REACTOME_TNFR2_NON_CANONICAL_NF_KB_PATHWAY<br>KEGG_T_CELL_RECEPTOR_SIGNALING_PATHWAY | CD27,CD40LG,PLCG1 |
|  | 7_tr | BIOCARTA_ARF_PATHWAY<br>REACTOME_GPVI_MEDIATED_ACTIVATION_CASCADE | PIK3CA,PIK3CG |
| metapathway_4 | 3_pr | REACTOME_INTERLEUKIN_10_SIGNALING<br>WP_OVERVIEW_OF_PROINFLAMMATORY_AND_PROFIBROTIC_MEDIATORS | CXCL10,TNFRSF1B,CXCL11,MMP9 |

**Table 6.** Metapathway analysis.

| metagen<br>e_2nd_<br>level | metage<br>ne_1st_<br>level | Enrichment result | Signatures |
| --- | --- | --- | --- |
| metapat<br>hway_6 | 2_hf | KEGG_OXIDATIVE_PHOSPHORYLATION<br>KEGG_PARKINSONS_DISEASE<br>WP_NONALCOHOLIC_FATTY_LIVER_DISEASE<br>KEGG_ALZHEIMERS_DISEASE<br>KEGG_HUNTINGTONS_DISEASE<br>KEGG_OXIDATIVE_PHOSPHORYLATION<br>REACTOME_RESPIRATORY_ELECTRON_TRANSPORT | COX7B,COX7C,COX17,NDUFA1,NDUF<br>A4,UQCRFS1,NDUFAB1,NDUFB1,NDU<br>FB2,NDUFB4,NDUFB10,COX5B,SDHB |
| metapat<br>hway_7 | 0_hf | KEGG_OXIDATIVE_PHOSPHORYLATION<br>KEGG_PARKINSONS_DISEASE<br>WP_NONALCOHOLIC_FATTY_LIVER_DISEASE<br>KEGG_ALZHEIMERS_DISEASE<br>KEGG_HUNTINGTONS_DISEASE<br>REACTOME_RESPIRATORY_ELECTRON_TRANSPORT | NDUFA13,CS,COX5A,SLC25A4,ETFA,E<br>TFB,COX7C,COX8A,UQCRC1,NDUFA4,<br>UQCRC2,UQCRFS1,IDH2,NDUFAB1,N<br>DUFB2,NDUFB4,NDUFB10,NDUFS3,ND<br>UFS5,GOT1,SDHB,NDUFS6,GOT2 |
| metapat<br>hway_1 | 6_op | BIOCARTA_AHSP_PATHWAY<br>WP_HEMATOPOIETIC_STEM_CELL_DIFFERENTIATION<br>REACTOME_CLASS_I_MHC_MEDIATED_ANTIGEN_PROCESSIN<br>G_PRESENTATION | AHSP,GATA1,RIOK3,PIM1,KCNH2,KLF<br>1,LYL1,MXI1,MKRN1,UBA52,UBB,FBXO<br>9,UBE2H,MRC2,FBXO7,PSMF1,NEDD4<br>L,SIAH2,RNF14,RNF123,UBE2O,CDC34<br>,RNF182 |
|  | 2_op | REACTOME_PLATELET_ACTIVATION_SIGNALING_AND_AGGRE<br>GATION<br>REACTOME_RESPONSE_TO_ELEVATED_PLATELET_CYTOSOL<br>IC_CA2<br>KEGG_FOCAL_ADHESION | PDGFA,ITGA2B,ITGB1,ITGB3,ITGB5,V<br>WF |
| metapat<br>hway_0 | 3_hf | REACTOME_RESPONSE_OF_EIF2AK4_GCN2_TO_AMINO_ACID<br>_DEFICIENCY<br>REACTOME_NONSENSE_MEDIATED_DECAY_NMD |  |
|  | 4_op | REACTOME_RESPONSE_OF_EIF2AK4_GCN2_TO_AMINO_ACID<br>_DEFICIENCY<br>REACTOME_SELENOAMINO_ACID_METABOLISM<br>REACTOME_NONSENSE_MEDIATED_DECAY_NMD |  |
| metapat<br>hway_9 | 4_op | REACTOME_RESPONSE_OF_EIF2AK4_GCN2_TO_AMINO_ACID<br>_DEFICIENCY<br>REACTOME_SELENOAMINO_ACID_METABOLISM<br>REACTOME_NONSENSE_MEDIATED_DECAY_NMD |  |
|  | 5_op | REACTOME_ALPHA_DEFENSINS | DEFA1,DEFA3,DEFA4 |
|  | 1_op | KEGG_NATURAL_KILLER_CELL_MEDIATED_CYTOTOXICITY<br>PID_IL12_2PATHWAY |  |
|  | 0_op | REACTOME_INTERFERON_ALPHA_BETA_SIGNALING<br>REACTOME_INTERFERON_GAMMA_SIGNALING |  |
|  | 6_op | BIOCARTA_AHSP_PATHWAY<br>WP_HEMATOPOIETIC_STEM_CELL_DIFFERENTIATION<br>REACTOME_CLASS_I_MHC_MEDIATED_ANTIGEN_PROCESSIN<br>G_PRESENTATION | AHSP,GATA1,RIOK3,PIM1,KCNH2,KLF<br>1,LYL1,MXI1,MKRN1,UBA52,UBB,FBXO<br>9,UBE2H,MRC2,FBXO7,PSMF1,NEDD4<br>L,SIAH2,RNF14,RNF123,UBE2O,CDC34<br>,RNF182 |
|  | 3_hf | REACTOME_RESPONSE_OF_EIF2AK4_GCN2_TO_AMINO_ACID<br>_DEFICIENCY<br>REACTOME_NONSENSE_MEDIATED_DECAY_NMD |  |
|  | 6_hf | REACTOME_RESPONSE_OF_EIF2AK4_GCN2_TO_AMINO_ACID<br>_DEFICIENCY<br>REACTOME_NONSENSE_MEDIATED_DECAY_NMD<br>REACTOME_SRP_DEPENDENT_COTRANSLATIONAL_PROTEIN<br>_TARGETING_TO_MEMBRANE<br>KEGG_MEDICUS_REFERENCE_TRANSLATION_INITIATION<br>WP_CYTOPLASMIC_RIBOSOMAL_PROTEINS<br>REACTOME_NONSENSE_MEDIATED_DECAY_NMD_INDEPEND<br>ENT_OF_THE_EXON_JUNCTION_COMPLEX_EJC |  |
|  | 5_hf | REACTOME_NONSENSE_MEDIATED_DECAY_NMD<br>WP_CYTOPLASMIC_RIBOSOMAL_PROTEINS<br>REACTOME_RESPONSE_OF_EIF2AK4_GCN2_TO_AMINO_ACID<br>_DEFICIENCY<br>REACTOME_NONSENSE_MEDIATED_DECAY_NMD_INDEPEND<br>ENT_OF_THE_EXON_JUNCTION_COMPLEX_EJC<br>REACTOME_BINDING_AND_UPTAKE_OF_LIGANDS_BY_SCAVE<br>NGER_RECEPTORS<br>REACTOME_SRP_DEPENDENT_COTRANSLATIONAL_PROTEIN<br>_TARGETING_TO_MEMBRANE |  |
|  | 2_hf | KEGG_OXIDATIVE_PHOSPHORYLATION<br>KEGG_PARKINSONS_DISEASE<br>WP_NONALCOHOLIC_FATTY_LIVER_DISEASE<br>KEGG_ALZHEIMERS_DISEASE<br>KEGG_HUNTINGTONS_DISEASE | COX7B,COX7C,COX17,NDUFA1,NDUF<br>A4,UQCRFS1,NDUFAB1,NDUFB1,NDU<br>FB2,NDUFB4,NDUFB10,COX5B,SDHB |
|  | 1_hf | REACTOME_NONSENSE_MEDIATED_DECAY_NMD<br>REACTOME_SELENOAMINO_ACID_METABOLISM<br>REACTOME_INFLUENZA_INFECTION | RPL23,RPL37A,RPS28,RPL41,RPS11 |
|  | 4_hf | REACTOME_INFLUENZA_INFECTION<br>REACTOME_NONSENSE_MEDIATED_DECAY_NMD | RPL23,RPS20,RPS26,RPL37A,RPS28,R<br>PLP2,RPL12,RPS8,RPS11,RPL18,RPL1<br>8A |
|  | 0_hf | KEGG_OXIDATIVE_PHOSPHORYLATION<br>KEGG_PARKINSONS_DISEASE<br>WP_NONALCOHOLIC_FATTY_LIVER_DISEASE<br>KEGG_ALZHEIMERS_DISEASE<br>KEGG_HUNTINGTONS_DISEASE |  |
| metapat<br>hway_8 | 8_op | WP_LEUKOTRIENE_METABOLIC_PATHWAY<br>WP_GLUCOCORTICOID_RECEPTOR_PATHWAY<br>WP_LEUKOTRIENE_METABOLIC_PATHWAYWP_VIWP_VITAMIN_D_RECEPTOR_PATHWAY |  |
|  | 7_op | KEGG_ALLOGRAFT_REJECTION |  |
|  | 9_op | KEGG_NOTCH_SIGNALING_PATHWAY | NOTCH1,NOTCH2,NUMBL,CIR1,CREB<br>BP,NCOR2,MAML1,EP300,MAML3 |
|  | 2_op | REACTOME_PLATELET_ACTIVATION_SIGNALING_AND_AGGREGATION<br>REACTOME_RESPONSE_TO_ELEVATED_PLATELET_CYTOSOLIC_CA2<br>KEGG_FOCAL_ADHESION | PDGFA,ITGA2B,ITGB1,ITGB3,ITGB5,V<br>WF |
| metapat<br>hway_5 | 4_hf | REACTOME_INFLUENZA_INFECTION<br>REACTOME_NONSENSE_MEDIATED_DECAY_NMD | RPL23,RPS20,RPS26,RPL37A,RPS28,R<br>PLP2,RPL12,RPS8,RPS11,RPL18,RPL1<br>8A |

As we see in Figure 7-E, metapathway_8 and metapathway_1 contribute the most pain to RA patients, protects the patients against from the damage of REACTOME_RESPONSE_TO_ELEVATED_PLATELET_CYTOSOLIC_CA2 while metapathway_9 is the most enriched for RA in Figure 7-D, protecting the patients from the damage of metapathway_6, metapathway_7 and metapathway_0.

In Figure 7-F, we find metapathway_5 is top one, ranking before metapathway_6 and metapathway_7. Therefore, we assumed metapathway_5 is also a buffer considering its negative relationship with metapathway_6 and metapathway_7 in Figure 7-C. We checked its enrichment result in Table 6 and found it is enriched in type I IFN as well as its Gene Ontology Molecular Function including “oxidoreduction-driven active transmembrane transporter activity” with signatures COX7B, COX8A, UQCRC1, NDUFS5, COX6B1, related to signatures in metapathway_6(Table 6). However, generally speaking, the negative regulation is not as good as RA under type II IFN because metapathway_6 and metapathway_7 are still top-listed in Figure 7-EF and metapathway_7 is still high expressed in Figure 7-D. Under the same logic, metapathway_2 and metapathway_3 are better severity buffers enriched in type I IFN but already overwhelmed in Figure 7-D. Furthermore, metapathway_2 (6_hf) is more aligned with IFN-I’s negative feedback mechanisms because it focuses on translation repression, stress responses, and inflammatory mRNA degradation, while metapathway_3 (5_hf) has additional immune activation components, such as binding and uptake of ligands by scavenger receptors. So, the difference of the two diseases is different metapathways regulate the damage of the tissue, and it seems that RA patients have more ways to buffer the damage.

Figure 7-G shows the interactions between metagenes, we can see the most frequent one (metagene_3_hf, metagene_4_op) represents for the interaction between Response of EIF2AK4 (GCN2) to Amino Acid Deficiency and Nonsense-Mediated Decay (NMD) in both diseases^60^.

## Conclusion

based on our framework, we found difference in the similarity of the two totally different diseases RA and HF. We found type I IFN alone can also call a negative feedback loop and act as a buffer for disease severity, especially for HF, but not as good as type IL-10 or type II IFN for RA^61,62^.

### Case Study 6: Deciphering Pre-Ciliated Cell Plasticity and Multicellular Stress Response Under RVCSE Exposure

In this case we used the case from Causal identification of single-cell experimental perturbation effects with CINEMA-OT^18^ for single cell illustration. In the original paper, CINEMA-OT applies ICA in the context of cell types, with the goal of identifying cell-type-specific signals from complex multi-omics data. This is the reason why the author mentioned at the same time that the default setting has low tolerance for ICA convergence, which harms CINEMA-OT little in practice^63^. Besides, in the original paper the authors found Pre-ciliated cells co-exposed to both viral infection and CSE showed synergistic induction of genes encoding secreted proteins that are typically associated with secretory cells in resting cultures, including SCGB3A1, LCN2, BPIFB1, SLPI, and WFDC2. However, they didn’t further explore the mechanism and its interaction with other cell types.

We did metapathway analysis combining cell types and 1^st^ level cohort-ICA processed data with 20 components for 4 different batches: MOCK, RV, CSE and RVCSE. Currently, RNAcompare only supports 2 batches, so we did this case manually. In our use case, we used ICA to decompose components without thinking cell types at first. Then we introduced the 2^nd^ level ICA, combining the output with cell types together, treating cell types as one interaction term with metagenes. Through this way, we bypassed the limitation and addressed the question mentioned above.

The first 5 metapathways are shown in Figure 8-A. We can see for example metapathway_0 includes the interactions among Doublets, Pre-ciliated, Ionocyte and Brush+PNEC cells with their communications via related pathways. Figure 8-B is just the cell types we extracted from the metapathways, based on which, we drew the inter-cell communication plot in Figure 8-C. The size of the cell types can basically represent the synergy score: cell_type_Pre-ciliated(9) means Pre-ciliated cells exist in 9 metapathways. Therefore, we can easily identify the hub cell type(s): Brush+PNEC(11), Ionocyte(9) and Pre-ciliated(9) based on 4 batches. We then drew the Pearson Correlation matrix of metapathways in Figure 8-D for later use.

**Figure 8.**
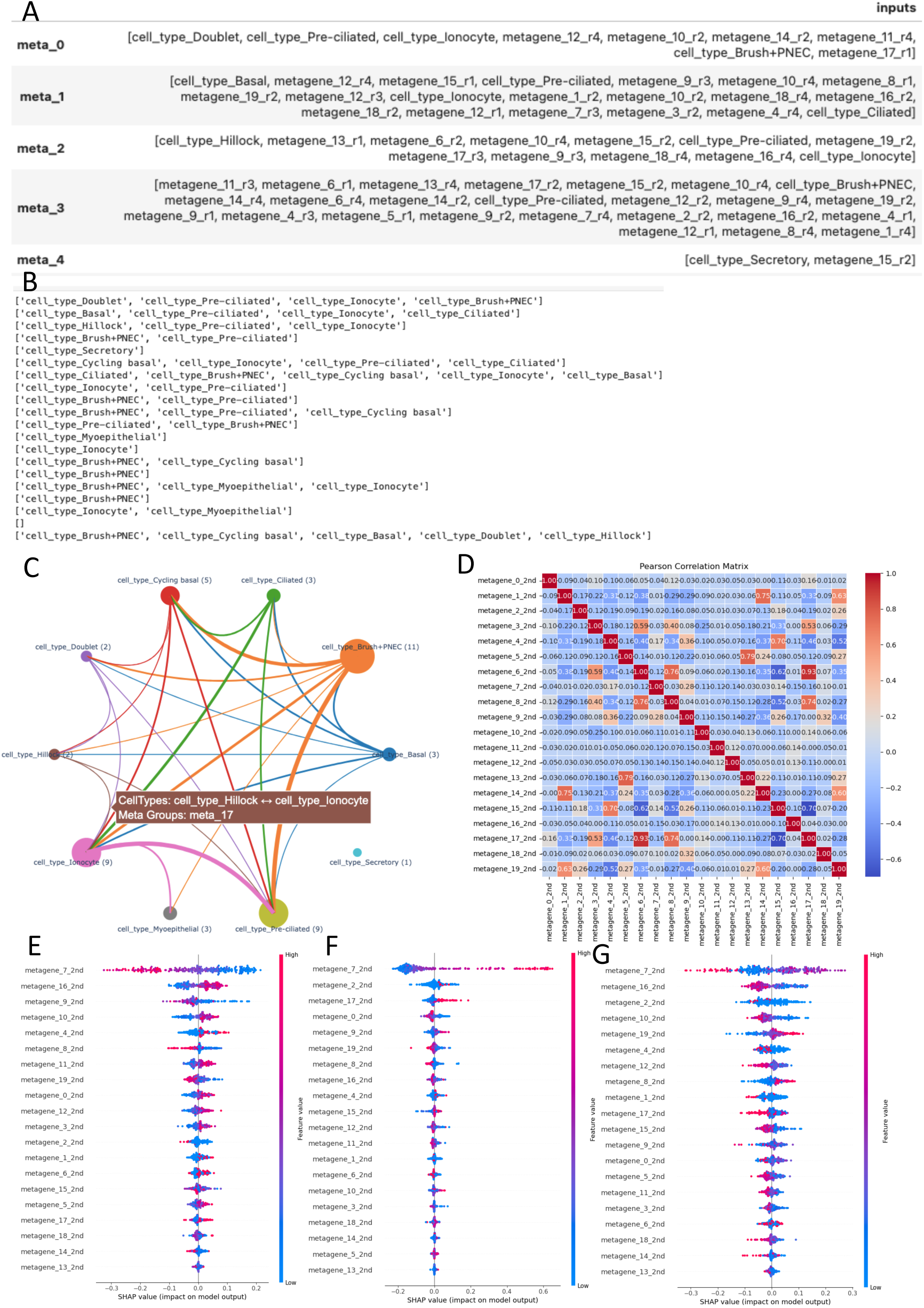
(A) Components of metapathway analysis; (B) Components of metapathway analysis for only cell types; (C) Relationship of cell types; (D) Pearson Correlation Matrix of metapathways; (E) SHAP importance plot for CSE vs others; (F) SHAP importance plot for RV vs others; (G) SHAP importance plot for RVCSE vs others;

We manually trained the model based on the 4 batches together. Figure 8-E-G are SHAP importance plot for CSE(labelled 1) vs others(labelled 0), RV(labelled 1) vs others(labelled 0), RVCSE(labelled 1) vs others(labelled 0) based on 1% test data.

After comparision of the 3 plots, we can easily see metapathway_9 and metapathway_17 are suppressed while metapathway_12 is promoted under RVCSE vs others. metapathway_9 is the interaction among Brush+PNEC, Pre-ciliated and Cycling basal; metapathway_17 is the interaction between Ionocyte and Pre-ciliated; metapathway_12 is Ionocyte. We then checked them with Figure 8-E-G and found metapathway_9 is suppressed by metapathway_19; metapathway_17 is suppressed by metapathway_4. Since the original paper only mentioned synergy effect with Pre-ciliated under RVCSE, we only analyse metapathway_19 here.

Table 7 shows the enrichment results for the related metapathways:

**Table 7.**
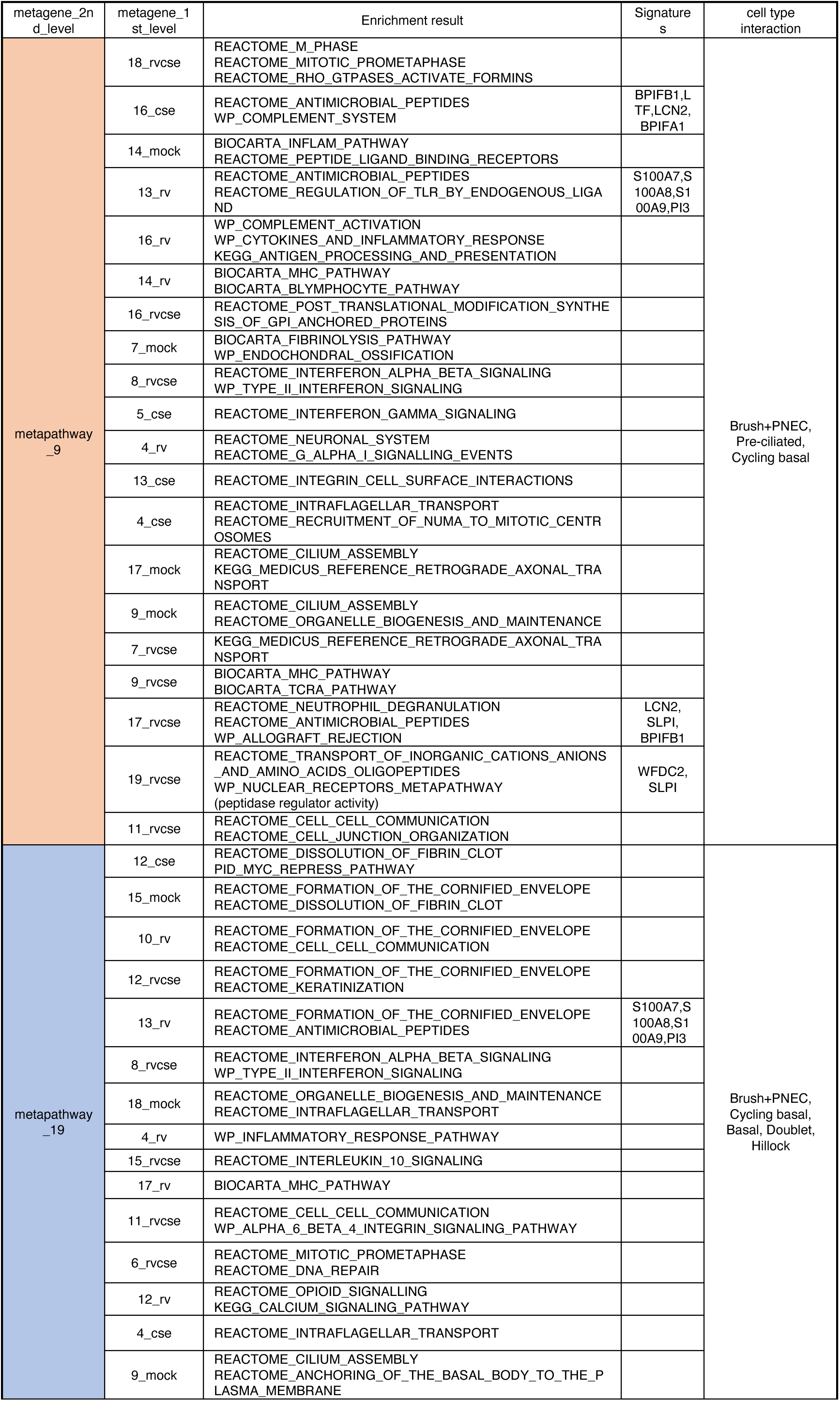
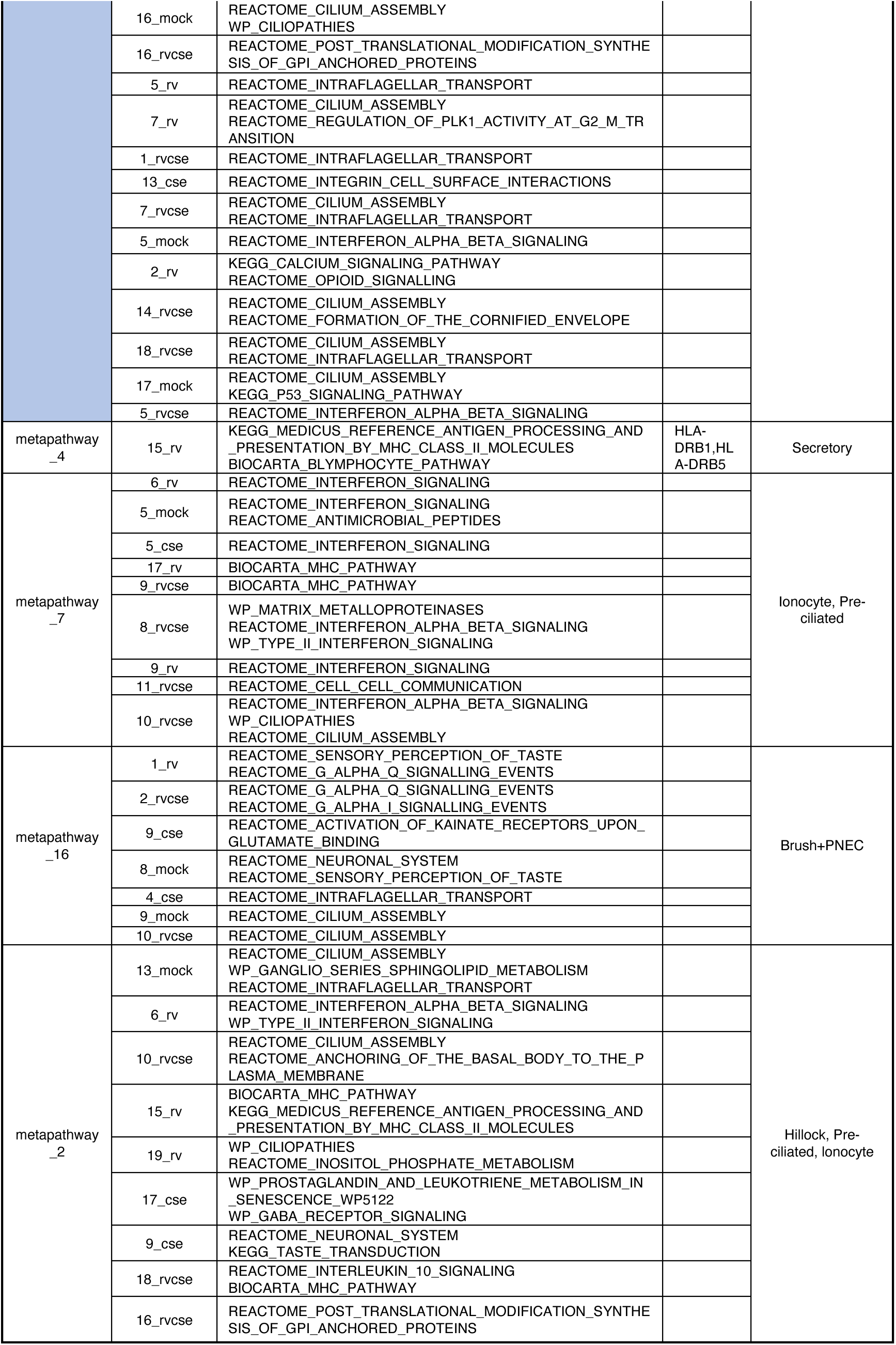
Metapathway analysis.

#### Metapathway Feedback Loop

Metapathway_9 includes IFN signaling and complement activation, forming a regulatory feedback loop during inflammatory stimuli; It also includes metagene_16_cse, which is defined by antimicrobial peptide (AMP) signatures (BPIFB1, LCN2), consistent with innate immune activation in the original paper.

#### Pre-Ciliated Cell Plasticity Under Stress (RVCSE)

Under combined respiratory virus (RV) and cigarette smoke extract (CSE) exposure, pre-ciliated cells undergo functional reprogramming:

Phase 1: Their secretion of inflammation-induced AMPs (metagene_13_rv) is suppressed by the proliferation process from metapathway_19.

Phase 2: They shift toward producing secretory granule lumen proteins (BPIFA1^64^, LTF^65^, LCN2^66^), which mediate intercellular communication characterised by the pathway REACTOME_CELL_CELL_COMMUNICATION. This secretory transition activates neighboring secretory cells, driving enhanced antigen processing and T cell activation to bring innate and adaptive immunity.

#### Multicellular Stress Response Orchestration

Pre-ciliated cells further recruit Basal, Doublet, and Hillock cells to amplify antimicrobial defences. These cells secrete proliferative AMPs (S100A7, S100A8, S100A9, PI3), which (1) directly neutralize pathogens (e.g., via S100A proteins’ metal chelation^67^); (2) amplify epithelial stress signaling (e.g., PI3’s protease inhibition^68^).

Besides, we also checked metapathway_7, enriched in IFN signalling, aligning with the description in the original paper under RV.

### Case Study 7: Cross-Species Metapathway Crosstalk Identifies Keratinization as a Novel Biomarker for Malaria Susceptibility

In this case, we combined GSE288146 and GSE289197 to explore the crosstalk of metapathways for host-parasite interaction where we used customised pipeline of RNAchat.

We first manually did the 1^st^ level ICA decomposition on the two datasets separately (10 components each) and combined them with an outer join. Then we filled the missing values with 0 for the two species after outer join.

Through metapathway analysis of both human and Plasmodium-derived pathways, we identified two major mechanisms by which the parasite interacts with host: metapathway_0 and metapathway_4, representing 2 different stages of infection: (Immune Modulation and Metabolic Adaptation) and (Host-Cell Structure Remodeling and Direct Cellular Interference).

In Table 8, we can see metapathway_6, metapathway_3 and metapathway_5 are the downstream stage of malaria infection. metapathway_1 is the immune response.

**Table 8.** Metapathway analysis.

| metagene_2nd_level | metagene_1st_level | Enrichment result |
| --- | --- | --- |
| metapathway_0 | 4_hum | REACTOME_GAP_JUNCTION_TRAFFICKING_AND_REGULATION |
|  | 9_hum | icosanoid transport<br>regulation of fatty acid transport<br>cellular response to steroid hormone stimulus |
|  | 0_hum | REACTOME_NEUTROPHIL_DEGRANULATION<br>REACTOME_SRP_DEPENDENT_COTRANSLATIONAL_PROTEIN_TARGETING_TO_MEMBRANE<br>REACTOME_EUKARYOTIC_TRANSLATION_INITIATION |
|  | 4_plas | structural constituent of chromatin<br>protein dimerization activity |
|  | 2_hum | adaptive thermogenesis<br>regulation of peptide hormone secretion |
|  | 3_plas | structural constituent of ribosome<br>structural molecule activity<br>RNA binding<br>Peptide chain elongation<br>Selenoamino acid metabolism<br>Nonsense-Mediated Decay (NMD)<br>Eukaryotic Translation Initiation |
|  | 6_plas | Cellular response to heat stress<br>Regulation of HSF1-mediated heat shock response<br>Protein processing in endoplasmic reticulum |
|  | 8_plas | PKMTs methylate histone lysines |
| metapathway_4 | 0_hum | REACTOME_NEUTROPHIL_DEGRANULATION<br>REACTOME_SRP_DEPENDENT_COTRANSLATIONAL_PROTEIN_TARGETING_TO_MEMBRANE<br>REACTOME_EUKARYOTIC_TRANSLATION_INITIATION |
|  | 1_hum | cadherin binding<br>exonuclease activity |
|  | 5_hum | heart looping<br>erythrocyte differentiation<br>myeloid cell homeostasis |
|  | 2_plas | protein dimerization activity<br>symbiont-containing vacuole membrane<br>extracellular membrane-bounded organelle |
|  | 0_plas | host cell cytoplasm<br>host intracellular part |
|  | 4_hum | REACTOME_GAP_JUNCTION_TRAFFICKING_AND_REGULATION |
|  | 9_plas | extracellular region |
| metapathway_2 | 3_hum | oxygen binding |
| metapathway_1 | 6_hum | REACTOME_NEUTROPHIL_DEGRANULATION<br>REACTOME_NONSENSE_MEDIATED_DECAY_NMD |
| metapathway_6 | 1_hum | cadherin binding<br>exonuclease activity |
|  | 9_hum | icosanoid transport<br>regulation of fatty acid transport |
|  | 8_hum | REACTOME_KERATINIZATION<br>REACTOME_FORMATION_OF_THE_CORNFIED_ENVELOPE |
|  | 5_hum | erythrocyte differentiation<br>myeloid cell homeostasis |
|  | 4_hum | fatty acid metabolic process<br>erythrocyte differentiation |
| metapathway_3 | 2_hum | adaptive thermogenesis<br>regulation of peptide hormone secretion |
|  | 4_hum | REACTOME_GAP_JUNCTION_TRAFFICKING_AND_REGULATION |
|  | 5_hum | heart looping<br>erythrocyte differentiation<br>myeloid cell homeostasis |
|  | 8_hum | REACTOME_KERATINIZATION<br>REACTOME_FORMATION_OF_THE_CORNFIED_ENVELOPE |
| metapathway_5 | 7_hum | WP_IL3_SIGNALING_PATHWAY<br>WP_IL5_SIGNALING_PATHWAY<br>WP_VEGFAVEGFR2_SIGNAL |
|  | 5_hum | heart looping<br>erythrocyte differentiation |
|  | 9_hum | icosanoid transport<br>regulation of fatty acid transport |
|  | 4_hum | REACTOME_GAP_JUNCTION_TRAFFICKING_AND_REGULATION |

We then extracted the patients’ data alone and did Random Forests (0.1 test split) to their number of parasitemia (log1p-ed) in the blood (Figure 9-C) with test MSE 2.4652. As we see metapathway_0 and metapathway_4 present a complementary mechanism to keep the number of parasitemia at a fixed level while metapathway_6 took the dominant. Currently there are many papers trying to cure malaria with solutions related to metapathway_0 and metapathway_4^69–71^. However, no paper talks about metapathway_6 related to keratinization. Paper found that serum lipids can regulate keratinization^72–74^. Therefore, we think keratinization level reflects malaria susceptibility in terms of the following genes related to lipid metabolism: LIPM^75^, LIPK^76^, LIPN^77^.

**Figure 9.**
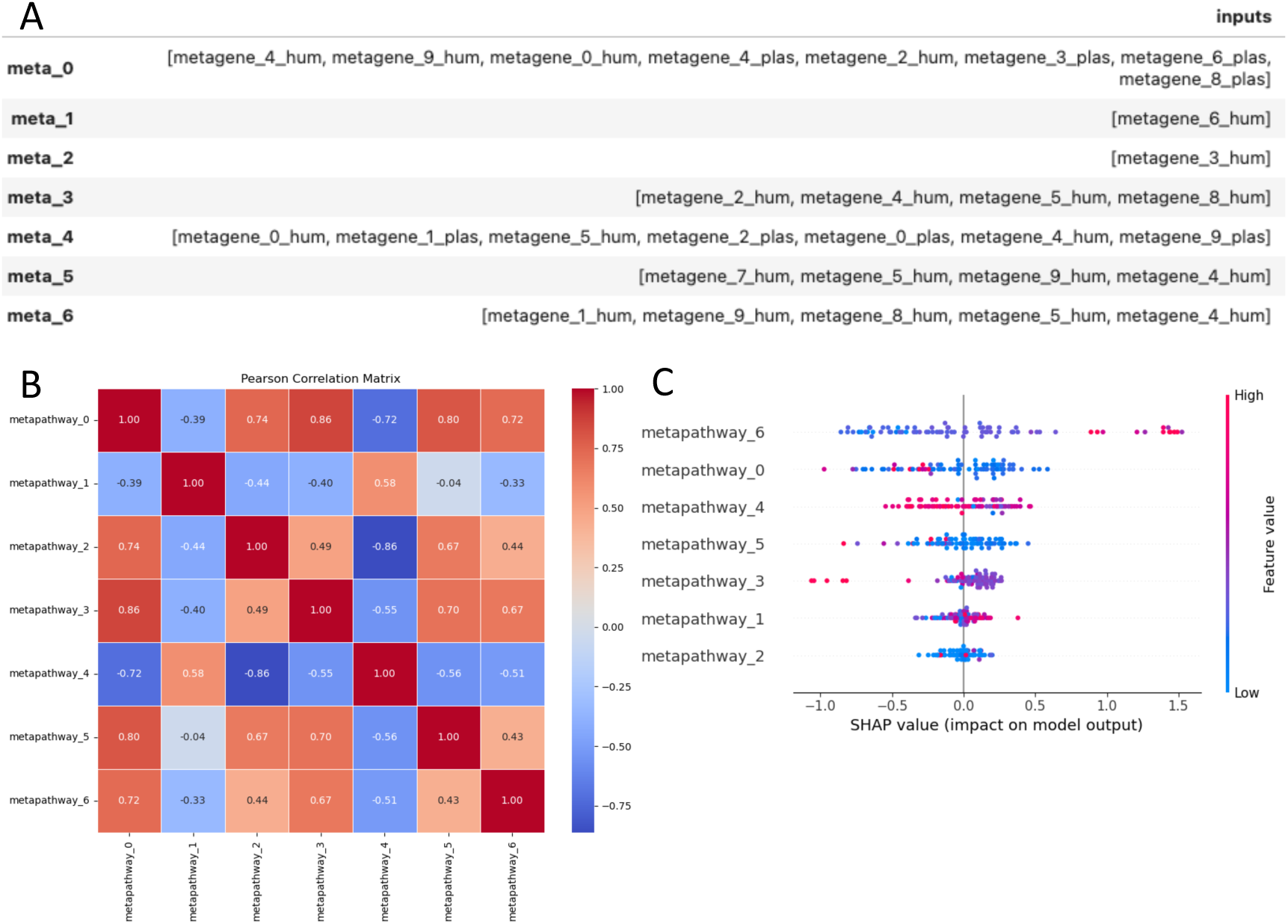
(A) Components of metapathway analysis; (B) Pearson Correlation Matrix of metapathways; (C) SHAP importance for metapathways (human only) to the number of parasitemia in the blood.

Appendix II shows the communications between metagenes.

## Discussion

RNAchat represents a paradigm shift in analyzing pathway and cell-cell communication by integrating clinical heterogeneity with multi-omics data through machine learning. Unlike existing tools such as CellChat or MetaPathways, which focus narrowly on predefined ligand-receptor interactions or microbial metabolic pathways, RNAchat prioritizes identifying clinically actionable metapathways—cooperative molecular modules that bridge omics layers and correlate with patient outcomes. Here, we discuss how RNAchat addresses critical gaps in the field, its biological and clinical implications, and its limitations.

### Clinical Integration as a Key Advancement

Most pathway analysis tools operate in a vacuum, dissecting molecular interactions without linking them to clinical phenotypes. RNAchat closes this gap by combining unsupervised ICA decomposition with supervised ML models to directly associate metapathways with outcomes like drug resistance, pain severity, or disease progression. By anchoring findings in clinical data, RNAchat transforms pathway analysis from a descriptive exercise into a predictive and actionable framework.

### Overcoming Limitations of Direction-Centric Tools

A key distinction between RNAchat and tools like CellChat is the lack of explicit directionality in inferred interactions. While CellChat prioritizes directional ligand-receptor signaling, it relies on curated databases that omit non-canonical interactions and cannot prioritize pathways based on functional outcomes. RNAchat circumvents these limitations by focusing on cooperative metapathways—groups of co-regulated genes that jointly influence clinical phenotypes.

### Biological Discoveries Enabled by RNAchat

#### 1. Therapy-Specific Metapathways

RNAchat uncovered that the same metagene (e.g., IFN-related *metagene*) participates in divergent metapathways depending on treatment context.

#### 2. Tissue-Specific Pathway Regulation

By comparing synovium and blood, RNAchat revealed that pain-associated feedback loops are negatively buffered in blood but dysregulated in synovial tissue. This suggests that systemic biomarkers may poorly reflect local disease mechanisms, advocating for tissue-targeted therapies.

#### 3. Conserved Stress-Response Strategies of HF

RNAchat identified IFN signaling as a universal stress-response hub across RA and HF, but with disease-specific buffering strategies.

### Technical Innovations and Limitations

#### 1. Strengths

##### Hypothesis-Free Discovery

RNAchat’s ICA-based approach avoids biases from pre-defined databases, enabling identification of novel interactions (e.g., EJC-independent NMD and BCR signaling crosstalk).

##### Bulk and Single-Cell Compatibility

Unlike CellChat, RNAchat operates on bulk and single-cell data, distinguishing paracrine/autocrine signaling in bulk via differential pathway enrichment (e.g., synovium vs. blood).

##### Explainable ML

SHAP values provide intuitive prioritization of metapathways.

#### 2. Limitations

##### Directionality

While RNAchat identifies cooperative metapathways, it cannot infer directional cell-cell communication. We propose combining RNAchat with CellChat in single-cell datasets to resolve this.

##### Validation Dependency

RNAchat’s predictions require external experimental validation, though we mitigated this by cross-referencing high-impact studies.

### Future Directions

RNAchat’s modular design allows expansion into new domains:

#### Non-Immune Diseases

Applying RNAchat to cancer or neurodegeneration could reveal conserved stress-response metapathways.

#### Therapeutic Discovery

drug targeting for the prioritised metapathways.

#### Dynamic Modeling

Integrating time-series data to track metapathway evolution during treatment.

### Conclusion

RNAchat redefines pathway analysis by placing clinical heterogeneity at the core of multi-omics integration. Its ability to identify therapy-specific metapathways and resolve feedback mechanisms like the “pain buffer” provides a roadmap for personalized medicine. While tools like CellChat remain valuable for directional signaling, RNAchat addresses a critical unmet need: translating pathway crosstalk into clinically actionable insights. As precision medicine advances, RNAchat’s framework will prove indispensable for dissecting the molecular basis of treatment response and disease heterogeneity.

## Supporting information

image and supplemental files

## Declarations

### Ethics approval and consent to participate

For the ORBIT data, all participants provided written, informed consent. The study protocol ORBIT ^14^ was approved by the West of Scotland Research Ethics Committee on 3/11/2009 (REC reference number: 09/S0703/109/ EudraCT number: 2009-011268-13)

For other studies, data were published.

### Consent for publication

We have consent for publication.

### Availability of data and materials

The code of can be found https://github.com/tangmingcan/RNAchat.

The RNA-Seq data from ORBIT can be found under the accession numbers Pending waiting for the publication of RNAcare, which is currently under review by BMC Medicine.

### Authors’ contributions

MT – Construction of tool. Data analysis. Writing of first draft.

## Acknowledgements

Mingcan previously worked at the School of Infection & Immunity, University of Glasgow, UK, where this study originated from his prior research with others. We also thank Senior Lecturer Tim Storer at computing science, University of Glassgow, Professor James Pan and Jan Chow from the Business Analytics Department at the National University of Singapore for their helps to this paper.

## Appendix I

In this section, we added HSP90AA1 to the 1^st^ level ICA result, then did 2^nd^ level ICA again. This is another analysis, so the metagenes are not exactly matched with the ones with the same numbers in the case study. Figure 1 shows its interaction with other megagenes under three situations.

Figure 1-A includes two drugs and we can see HSP90AA1 interacted with metagene_3_1st, which is IFN signalling in Figure 1-D; Figure 1-B includes only anti-TNF and we can see HSP90AA1 interacted with metagene_7_1st, which is nonsense mediated decay, more related to type I IFN in Figure 1-E; Figure 1-C includes only Rituximab and we can see HSP90AA1 was more neutralised. Therefore, this can demonstrate its role change under different drugs.

Besides, we can check the interactions between L1CAM with other pathways., We first found L1CAM was metagene_2_1st (Figure 1-F), then we did its corresponding metapathway analysis in Table 1 based on all datasets, anti-TNF, and Rituximab respectively. From the table we can see, the common part is IFN singaling and L1CAM together proliferate the stimuli from platelet activation signaling aggregation and formation of fibrin clot clotting cascade. However, the introduction of Rituximab prioritises the order of type II IFN (not very obvious) while there is another interesting participant Rho GTPase Cycle (including HSP90AA1). It further proves that under this situation, Rho GTPase Cycle was released from interaction with B cell receptor signalling.

**Figure 1.**
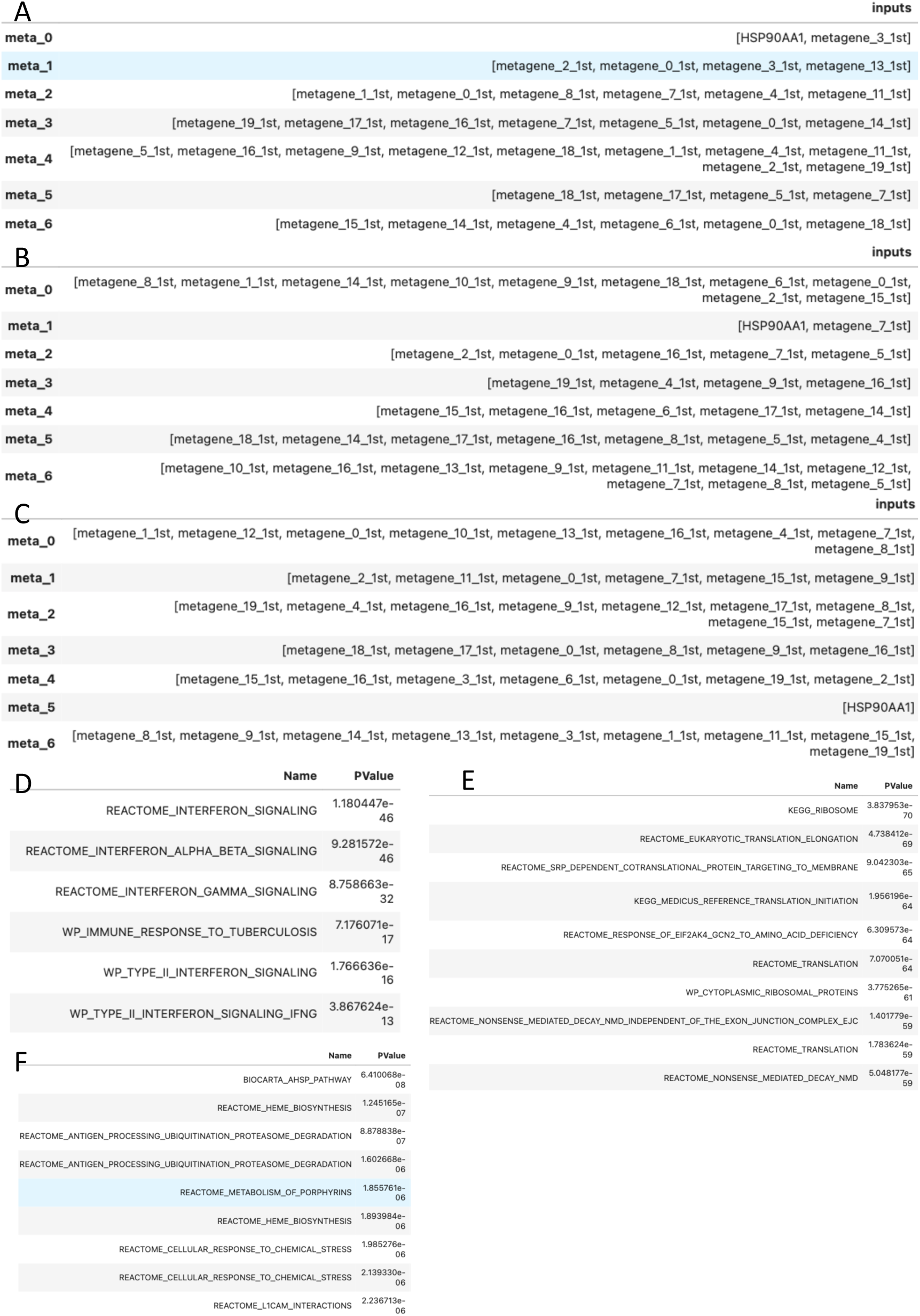
(A) Metapathway analysis for two drugs; (B) Metapathway analysis for Anti-TNF; (C) Metapathway analysis for Rituximab; (D) Enrichment analysis for metagene_3_1st; (E) Enrichment analysis for metagene_7_1st. (F) Enrichment analysis for metagene_2_1^st^.

**Table 1.** Metapathway analysis.

| metagene_2nd_level | metagene_1st_level | Enrichment reslut |
| --- | --- | --- |
| metapathway_1(All) | metagene_2_1st | REACTOME_L1CAM_INTERACTIONS |
|  | metagene_0_1st | REACTOME_PLATELET_ACTIVATION_SIGNALING_AND_AGGREGATION<br>REACTOME_FORMATION_OF_FIBRIN_CLOT_CLOTTING_CASCADE |
|  | metagene_3_1st | REACTOME_INTERFERON_ALPHA_BETA_SIGNALING<br>REACTOME_INTERFERON_GAMMA_SIGNALING |
|  | metagene_13_1st | REACTOME_ANTIMICROBIAL_PEPTIDES |
| metapathway_2(Anti- TNF) | metagene_2_1st | REACTOME_L1CAM_INTERACTIONS |
|  | metagene_0_1st | REACTOME_PLATELET_ACTIVATION_SIGNALING_AND_AGGREGATION<br>REACTOME_FORMATION_OF_FIBRIN_CLOT_CLOTTING_CASCADE |
|  | metagene_16_1st | KEGG_ALLOGRAFT_REJECTION |
|  | metagene_7_1st | REACTOME_NONSENSE_MEDIATED_DECAY_NMD |
|  | metagene_5_1st | REACTOME_FCGR_ACTIVATION |
| metapathway_1(Rituximab) | metagene_2_1st | REACTOME_L1CAM_INTERACTIONS |
|  | metagene_11_1st | REACTOME_INTERFERON_GAMMA_SIGNALING<br>KEGG_ALLOGRAFT_REJECTION |
|  | metagene_0_1st | REACTOME_PLATELET_ACTIVATION_SIGNALING_AND_AGGREGATION<br>REACTOME_FORMATION_OF_FIBRIN_CLOT_CLOTTING_CASCADE |
|  | metagene_7_1st | REACTOME_NONSENSE_MEDIATED_DECAY_NMD |
|  | metagene_15_1st | REACTOME_RHO_GTPASE_CYCLE |
|  | metagene_9_1st | PID_IL12_2PATHWAY<br>BIOCARTA_NO2IL12_PATHWAY |

## Appendix II

We can also plot the diagram based on metagenes colored by 4 batches with a designated threshold controlling the frequency of two interacted metagenes existing in the metapathways. As we see in the Figure 1, metagene_9_mock interacts with metagene_4_cse under CSE/MOCK; metagene_9_mock interacts with metagene_6_rvcse under RVCSE/MOCK and metagene_4_cse interacts with metagene_6_rvcse under RVCSE/CSE. Therefore, we can know the relationships (promotion/suppression) of the 3 metagenes by just further plotting a Pearson matrix plot of these metagenes.

**Figure 1.**
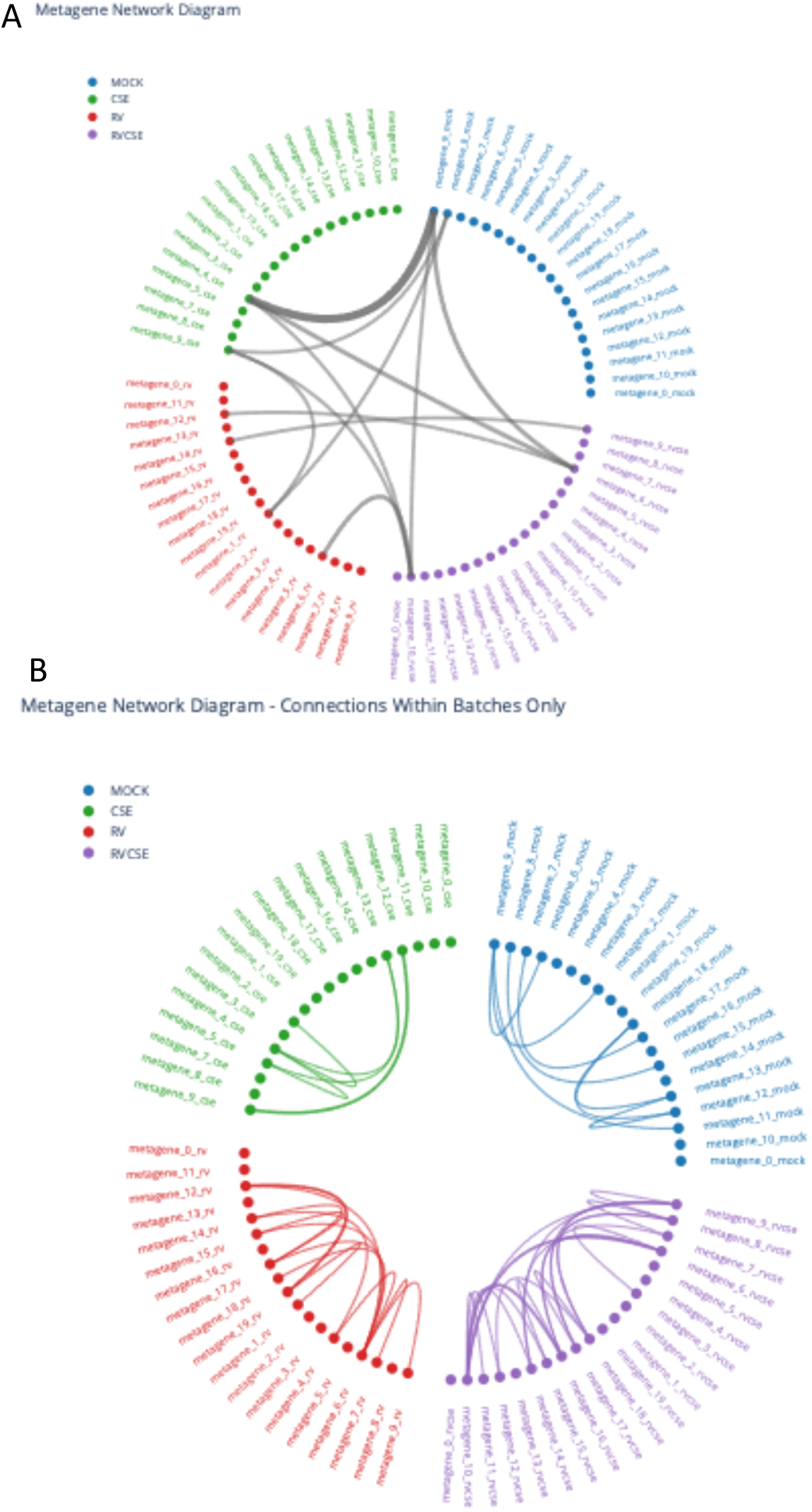
(A) Relationship of metagenes under different perturbations; (B) Relationship of metagenes under same perturbation.

