## Supplementary material for "RNAchat: Integrating machine learning algorithms to identify metapathways based on clinical and multi-omics data": image and supplemental files

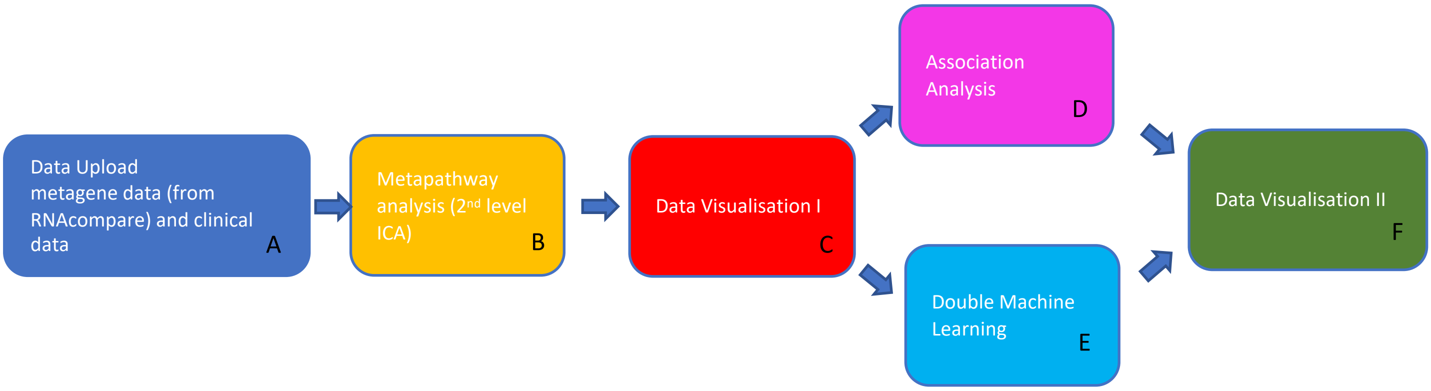
Data Upload
metagene data (from RNAcompare) and clinical data
A
Metapathway analysis (2nd level ICA)
B
Data Visualisation I
C
Association Analysis
D
Double Machine Learning
E
Data Visualisation II
F

A

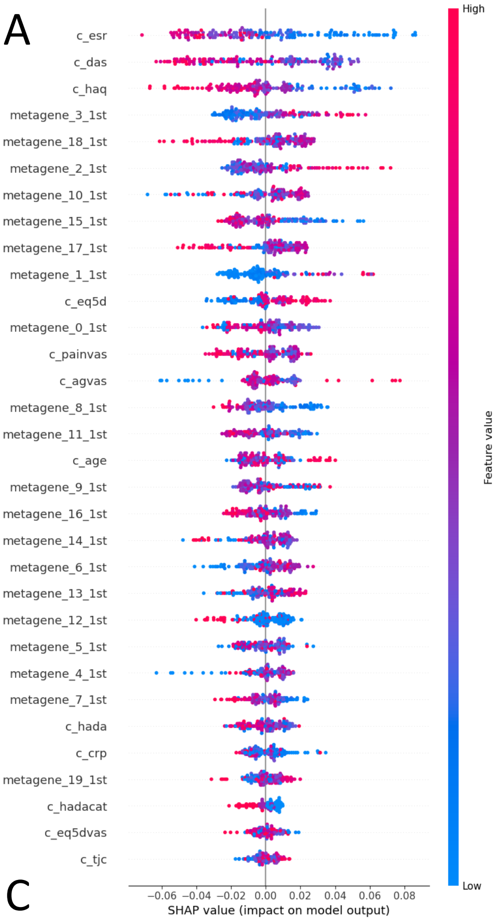


B

| Name | PValue |
|---|---|
| REACTOME_INTERFERON_SIGNALING | 1.476452e-46 |
| REACTOME_INTERFERON_ALPHA_BETA_SIGNALING | 1.071163e-45 |
| REACTOME_INTERFERON_GAMMA_SIGNALING | 9.851463e-32 |
| WP_IMMUNE_RESPONSE_TO_TUBERCULOSIS | 7.511500e-17 |
| WP_TYPE_II_INTERFERON_SIGNALING | 1.861285e-16 |
| WP_TYPE_II_INTERFERON_SIGNALING_IFNG | 4.031285e-13 |

C

| | inputs |
|---|---|
| **meta_0** | [metagene_1_1st, metagene_0_1st, metagene_10_1st, metagene_4_1st, metagene_7_1st] |
| **meta_1** | [metagene_2_1st, metagene_0_1st, metagene_13_1st, metagene_6_1st] |
| **meta_2** | [metagene_16_1st, metagene_14_1st, metagene_7_1st, metagene_3_1st, metagene_17_1st, metagene_8_1st, metagene_0_1st] |
| **meta_3** | [metagene_18_1st, metagene_17_1st, metagene_4_1st, metagene_8_1st, metagene_10_1st] |
| **meta_4** | [metagene_19_1st, metagene_17_1st, metagene_4_1st, metagene_14_1st, metagene_9_1st, metagene_8_1st] |
| **meta_5** | [metagene_8_1st, metagene_15_1st, metagene_6_1st, metagene_19_1st, metagene_7_1st] |
| **meta_6** | [metagene_6_1st, metagene_17_1st, metagene_15_1st, metagene_8_1st] |

D

| | inputs |
|---|---|
| **meta_0** | [metagene_2_1st, metagene_0_1st, metagene_14_1st, metagene_5_1st, metagene_6_1st] |
| **meta_1** | [metagene_12_1st, metagene_14_1st, metagene_11_1st, metagene_9_1st, metagene_13_1st, metagene_15_1st, metagene_10_1st, metagene_18_1st] |
| **meta_2** | [metagene_16_1st, metagene_14_1st, metagene_7_1st, metagene_17_1st, metagene_8_1st] |
| **meta_3** | [metagene_18_1st, metagene_17_1st, metagene_8_1st, metagene_4_1st] |
| **meta_4** | [metagene_8_1st, metagene_15_1st, metagene_6_1st, metagene_19_1st, metagene_7_1st, metagene_1_1st, metagene_18_1st] |
| **meta_5** | [metagene_19_1st, metagene_17_1st, metagene_14_1st, metagene_9_1st, metagene_4_1st, metagene_8_1st, metagene_15_1st, metagene_5_1st] |
| **meta_6** | [metagene_6_1st, metagene_17_1st, metagene_15_1st, metagene_8_1st, metagene_3_1st] |

E

| | inputs |
|---|---|
| **meta_0** | [metagene_1_1st, metagene_10_1st, metagene_0_1st, metagene_11_1st, metagene_12_1st, metagene_4_1st, metagene_14_1st, metagene_18_1st] |
| **meta_1** | [metagene_2_1st, metagene_13_1st, metagene_0_1st, metagene_16_1st, metagene_10_1st, metagene_6_1st] |
| **meta_2** | [metagene_18_1st, metagene_4_1st, metagene_17_1st, metagene_13_1st, metagene_8_1st, metagene_14_1st, metagene_7_1st, metagene_10_1st] |
| **meta_3** | [metagene_16_1st, metagene_14_1st, metagene_7_1st, metagene_8_1st, metagene_17_1st, metagene_3_1st, metagene_2_1st] |
| **meta_4** | [metagene_19_1st, metagene_17_1st, metagene_4_1st, metagene_9_1st, metagene_14_1st, metagene_8_1st] |
| **meta_5** | [metagene_8_1st, metagene_15_1st, metagene_9_1st, metagene_11_1st, metagene_19_1st, metagene_16_1st] |
| **meta_6** | [metagene_17_1st, metagene_6_1st, metagene_15_1st, metagene_5_1st] |

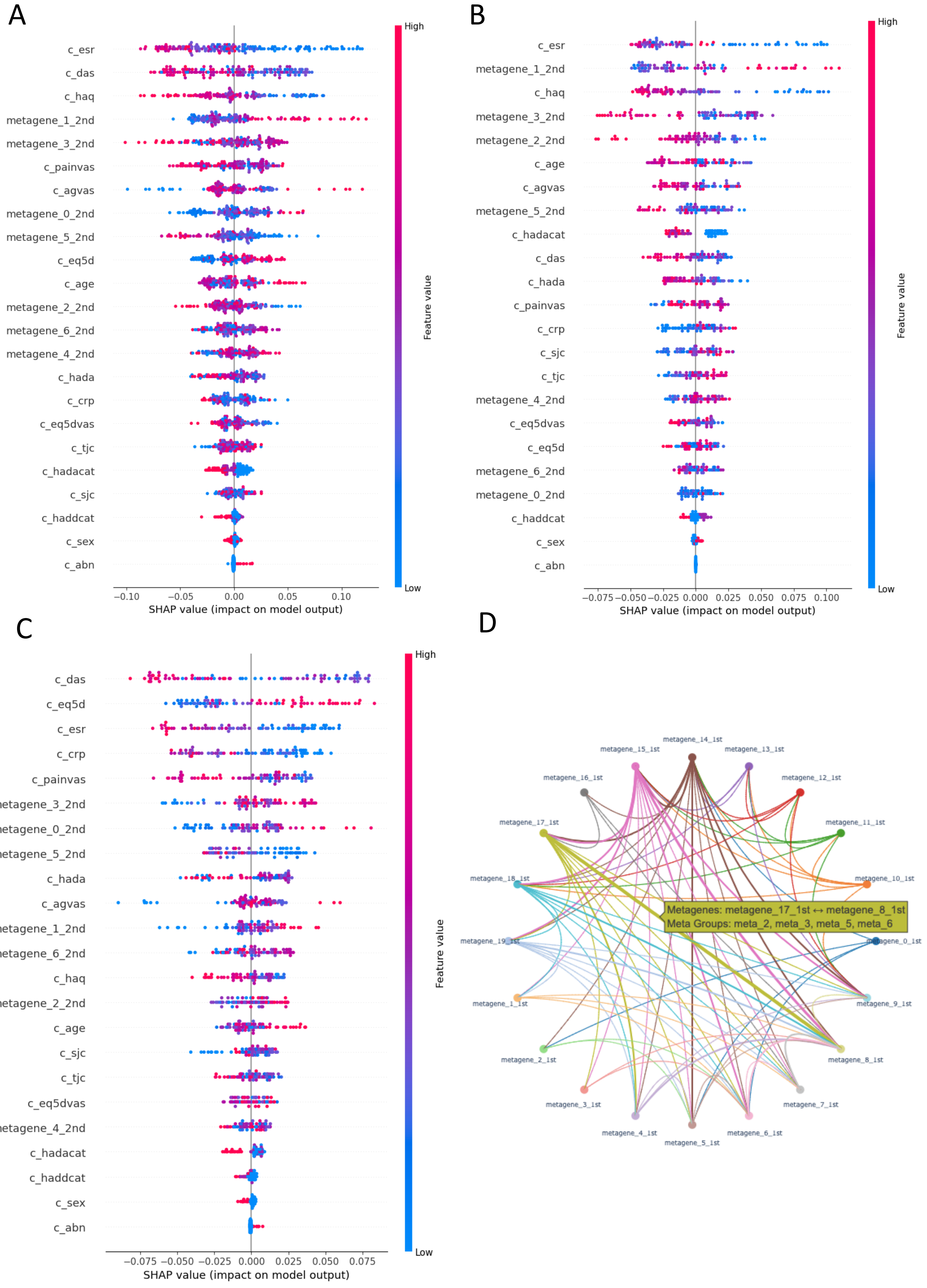
A
c_esr
c_das
c_haq
metagene_1_2nd
metagene_3_2nd
c_painvas
c_agvas
metagene_0_2nd
metagene_5_2nd
c_eq5d
c_age
metagene_2_2nd
metagene_6_2nd
metagene_4_2nd
c_hada
c_crp
c_eq5dvas
c_tjc
c_hadacat
c_sjc
c_haddcat
c_sex
c_abn
−0.10 −0.05 0.00 0.05 0.10
SHAP value (impact on model output)
High
Feature value
Low
B
c_esr
metagene_1_2nd
c_haq
metagene_3_2nd
metagene_2_2nd
c_age
c_agvas
metagene_5_2nd
c_hadacat
c_das
c_hada
c_painvas
c_crp
c_sjc
c_tjc
metagene_4_2nd
c_eq5dvas
c_eq5d
metagene_6_2nd
metagene_0_2nd
c_haddcat
c_sex
c_abn
−0.075 −0.050 −0.025 0.000 0.025 0.050 0.075 0.100
SHAP value (impact on model output)
High
Feature value
Low
C
c_das
c_eq5d
c_esr
c_crp
c_painvas
metagene_3_2nd
metagene_0_2nd
metagene_5_2nd
c_hada
c_agvas
metagene_1_2nd
metagene_6_2nd
c_haq
metagene_2_2nd
c_age
c_sjc
c_tjc
c_eq5dvas
metagene_4_2nd
c_hadacat
c_haddcat
c_sex
c_abn
−0.075 −0.050 −0.025 0.000 0.025 0.050 0.075
SHAP value (impact on model output)
High
Feature value
Low
D
metagene_14_1st
metagene_15_1st
metagene_13_1st
metagene_16_1st
metagene_12_1st
metagene_17_1st
metagene_11_1st
metagene_18_1st
metagene_10_1st
Metagenes: metagene_17_1st ↔ metagene_8_1st
Meta Groups: meta_2, meta_3, meta_5, meta_6
metagene_19_1st
metagene_0_1st
metagene_1_1st
metagene_9_1st
metagene_2_1st
metagene_8_1st
metagene_3_1st
metagene_7_1st
metagene_4_1st
metagene_6_1st
metagene_5_1st

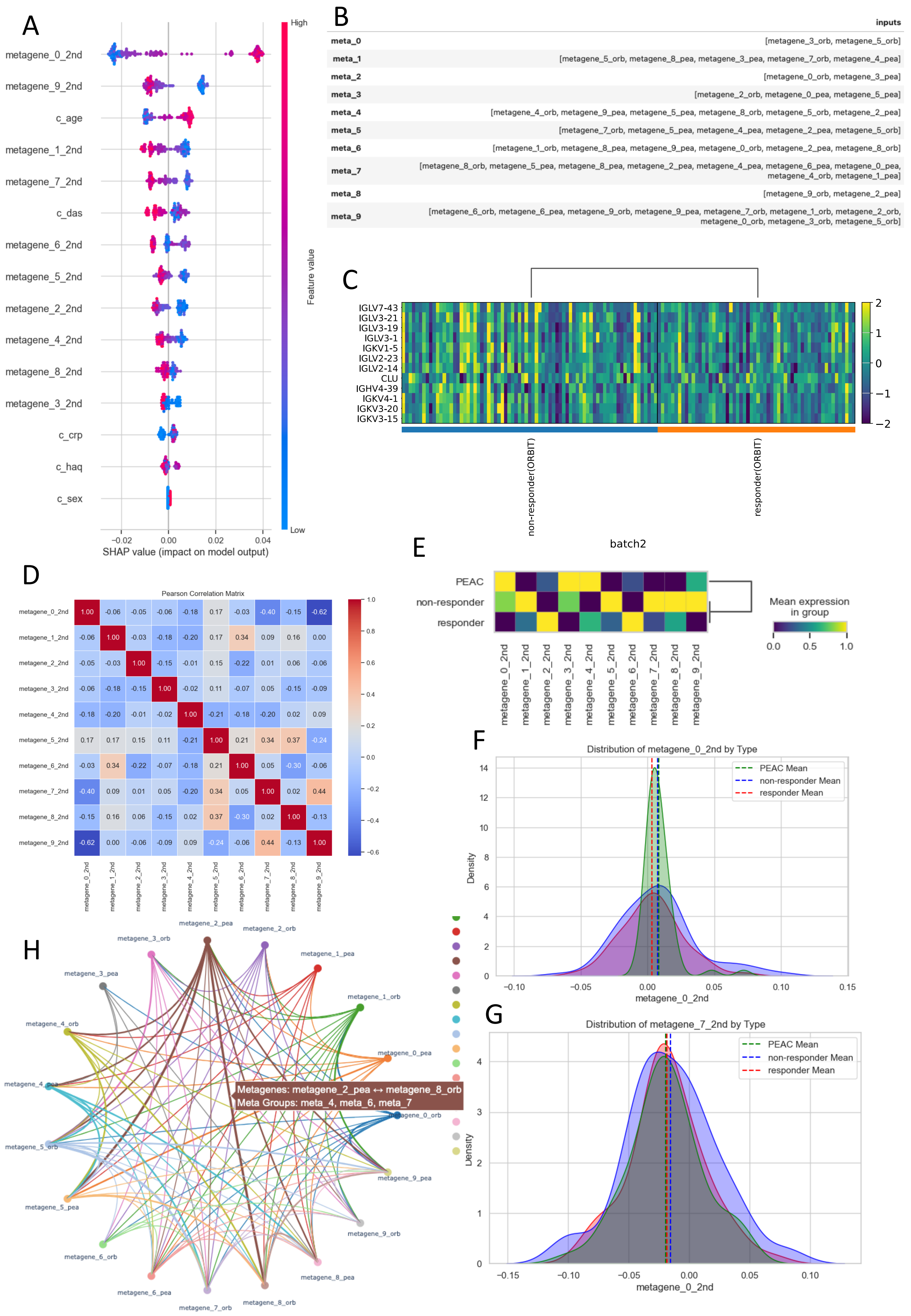

A
metagene_0_2nd
metagene_9_2nd
c_age
metagene_1_2nd
metagene_7_2nd
c_das
metagene_6_2nd
metagene_5_2nd
metagene_2_2nd
metagene_4_2nd
metagene_8_2nd
metagene_3_2nd
c_crp
c_haq
c_sex
−0.02 0.00 0.02 0.04
SHAP value (impact on model output)
High
Low
Feature value
B
inputs
meta_0 [metagene_3_orb, metagene_5_orb]
meta_1 [metagene_5_orb, metagene_8_pea, metagene_3_pea, metagene_7_orb, metagene_4_pea]
meta_2 [metagene_0_orb, metagene_3_pea]
meta_3 [metagene_2_orb, metagene_0_pea, metagene_5_pea]
meta_4 [metagene_4_orb, metagene_9_pea, metagene_5_pea, metagene_8_orb, metagene_5_orb, metagene_2_pea]
meta_5 [metagene_7_orb, metagene_5_pea, metagene_4_pea, metagene_2_pea, metagene_5_orb]
meta_6 [metagene_1_orb, metagene_8_pea, metagene_9_pea, metagene_0_orb, metagene_2_pea, metagene_8_orb]
meta_7 [metagene_8_orb, metagene_5_pea, metagene_8_pea, metagene_2_pea, metagene_4_pea, metagene_6_pea, metagene_0_pea, metagene_4_orb, metagene_1_pea]
meta_8 [metagene_9_orb, metagene_2_pea]
meta_9 [metagene_6_orb, metagene_6_pea, metagene_9_orb, metagene_9_pea, metagene_7_orb, metagene_1_orb, metagene_2_orb, metagene_0_orb, metagene_3_orb, metagene_5_orb]
C
IGLV7-43
IGLV3-21
IGLV3-19
IGLV3-1
IGKV1-5
IGLV2-23
IGLV2-14
CLU
IGHV4-39
IGKV4-1
IGKV3-20
IGKV3-15
non-responder(ORBIT)
responder(ORBIT)
2 1 0 −1 −2
D
Pearson Correlation Matrix
E
batch2
PEAC
non-responder
responder
Mean expression in group
0.0 0.5 1.0
F
Distribution of metagene_0_2nd by Type
PEAC Mean
non-responder Mean
responder Mean
Density
metagene_0_2nd
G
Distribution of metagene_7_2nd by Type
PEAC Mean
non-responder Mean
responder Mean
Density
metagene_0_2nd
H
Metagenes: metagene_2_pea ↔ metagene_8_orb
Meta Groups: meta_4, meta_6, meta_7

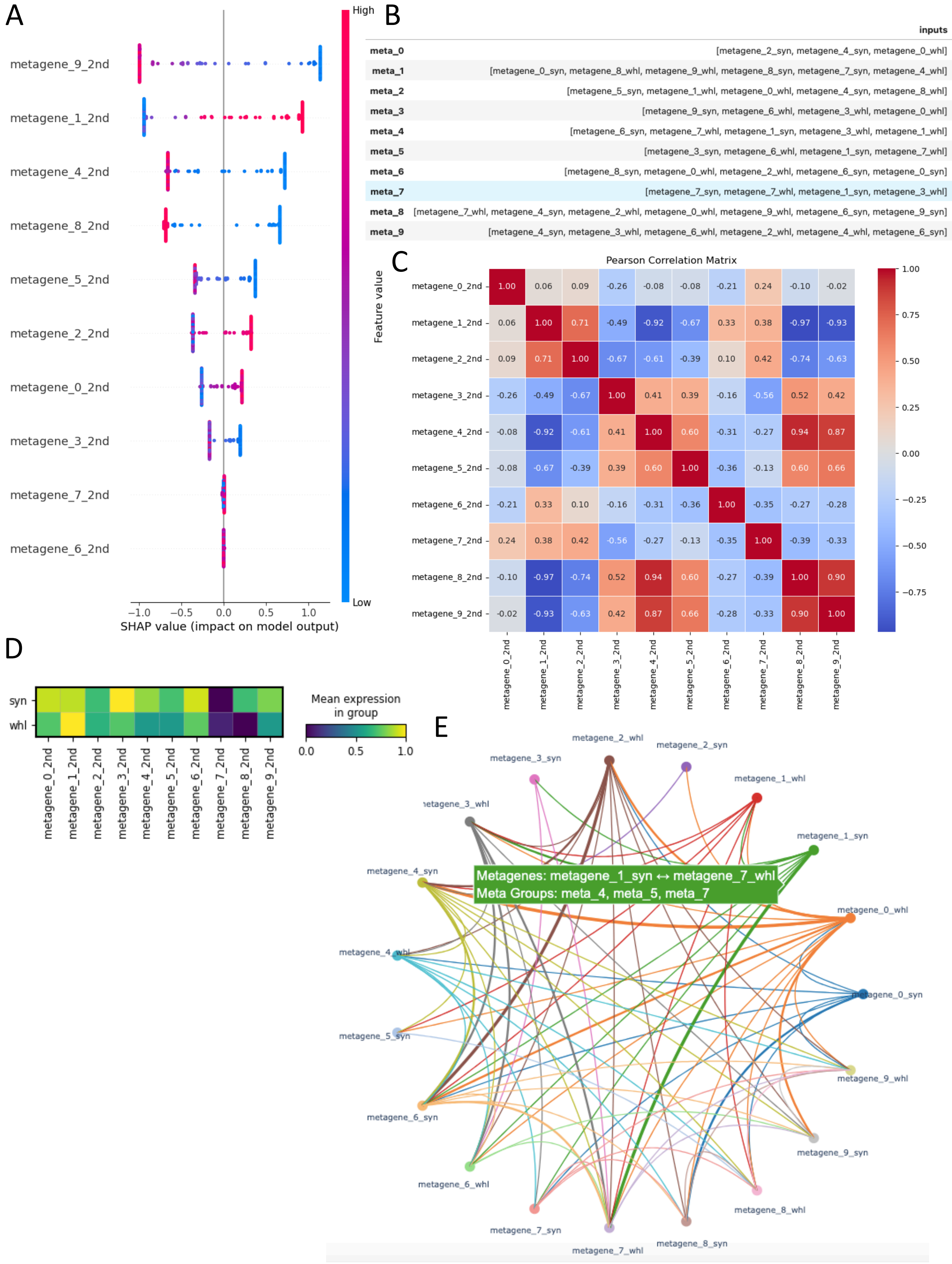
A
metagene_9_2nd
metagene_1_2nd
metagene_4_2nd
metagene_8_2nd
metagene_5_2nd
metagene_2_2nd
metagene_0_2nd
metagene_3_2nd
metagene_7_2nd
metagene_6_2nd
−1.0 −0.5 0.0 0.5 1.0
SHAP value (impact on model output)
High
Low
Feature value
B
inputs
meta_0 [metagene_2_syn, metagene_4_syn, metagene_0_whl]
meta_1 [metagene_0_syn, metagene_8_whl, metagene_9_whl, metagene_8_syn, metagene_7_syn, metagene_4_whl]
meta_2 [metagene_5_syn, metagene_1_whl, metagene_0_whl, metagene_4_syn, metagene_8_whl]
meta_3 [metagene_9_syn, metagene_6_whl, metagene_3_whl, metagene_0_whl]
meta_4 [metagene_6_syn, metagene_7_whl, metagene_1_syn, metagene_3_whl, metagene_1_whl]
meta_5 [metagene_3_syn, metagene_6_whl, metagene_1_syn, metagene_7_whl]
meta_6 [metagene_8_syn, metagene_0_whl, metagene_2_whl, metagene_6_syn, metagene_0_syn]
meta_7 [metagene_7_syn, metagene_7_whl, metagene_1_syn, metagene_3_whl]
meta_8 [metagene_7_whl, metagene_4_syn, metagene_2_whl, metagene_0_whl, metagene_9_whl, metagene_6_syn, metagene_9_syn]
meta_9 [metagene_4_syn, metagene_3_whl, metagene_6_whl, metagene_2_whl, metagene_4_whl, metagene_6_syn]
C
Pearson Correlation Matrix
metagene_0_2nd 1.00 0.06 0.09 -0.26 -0.08 -0.08 -0.21 0.24 -0.10 -0.02
metagene_1_2nd 0.06 1.00 0.71 -0.49 -0.92 -0.67 0.33 0.38 -0.97 -0.93
metagene_2_2nd 0.09 0.71 1.00 -0.67 -0.61 -0.39 0.10 0.42 -0.74 -0.63
metagene_3_2nd -0.26 -0.49 -0.67 1.00 0.41 0.39 -0.16 -0.56 0.52 0.42
metagene_4_2nd -0.08 -0.92 -0.61 0.41 1.00 0.60 -0.31 -0.27 0.94 0.87
metagene_5_2nd -0.08 -0.67 -0.39 0.39 0.60 1.00 -0.36 -0.13 0.60 0.66
metagene_6_2nd -0.21 0.33 0.10 -0.16 -0.31 -0.36 1.00 -0.35 -0.27 -0.28
metagene_7_2nd 0.24 0.38 0.42 -0.56 -0.27 -0.13 -0.35 1.00 -0.39 -0.33
metagene_8_2nd -0.10 -0.97 -0.74 0.52 0.94 0.60 -0.27 -0.39 1.00 0.90
metagene_9_2nd -0.02 -0.93 -0.63 0.42 0.87 0.66 -0.28 -0.33 0.90 1.00
1.00 0.75 0.50 0.25 0.00 −0.25 −0.50 −0.75
D
syn
whl
metagene_0_2nd metagene_1_2nd metagene_2_2nd metagene_3_2nd metagene_4_2nd metagene_5_2nd metagene_6_2nd metagene_7_2nd metagene_8_2nd metagene_9_2nd
Mean expression in group
0.0 0.5 1.0
E
Metagenes: metagene_1_syn ↔ metagene_7_whl
Meta Groups: meta_4, meta_5, meta_7
metagene_2_whl
metagene_2_syn
metagene_3_syn
metagene_1_whl
metagene_3_whl
metagene_1_syn
metagene_4_syn
metagene_0_whl
metagene_4_whl
metagene_0_syn
metagene_5_syn
metagene_9_whl
metagene_6_syn
metagene_9_syn
metagene_6_whl
metagene_8_whl
metagene_7_syn
metagene_8_syn
metagene_7_whl

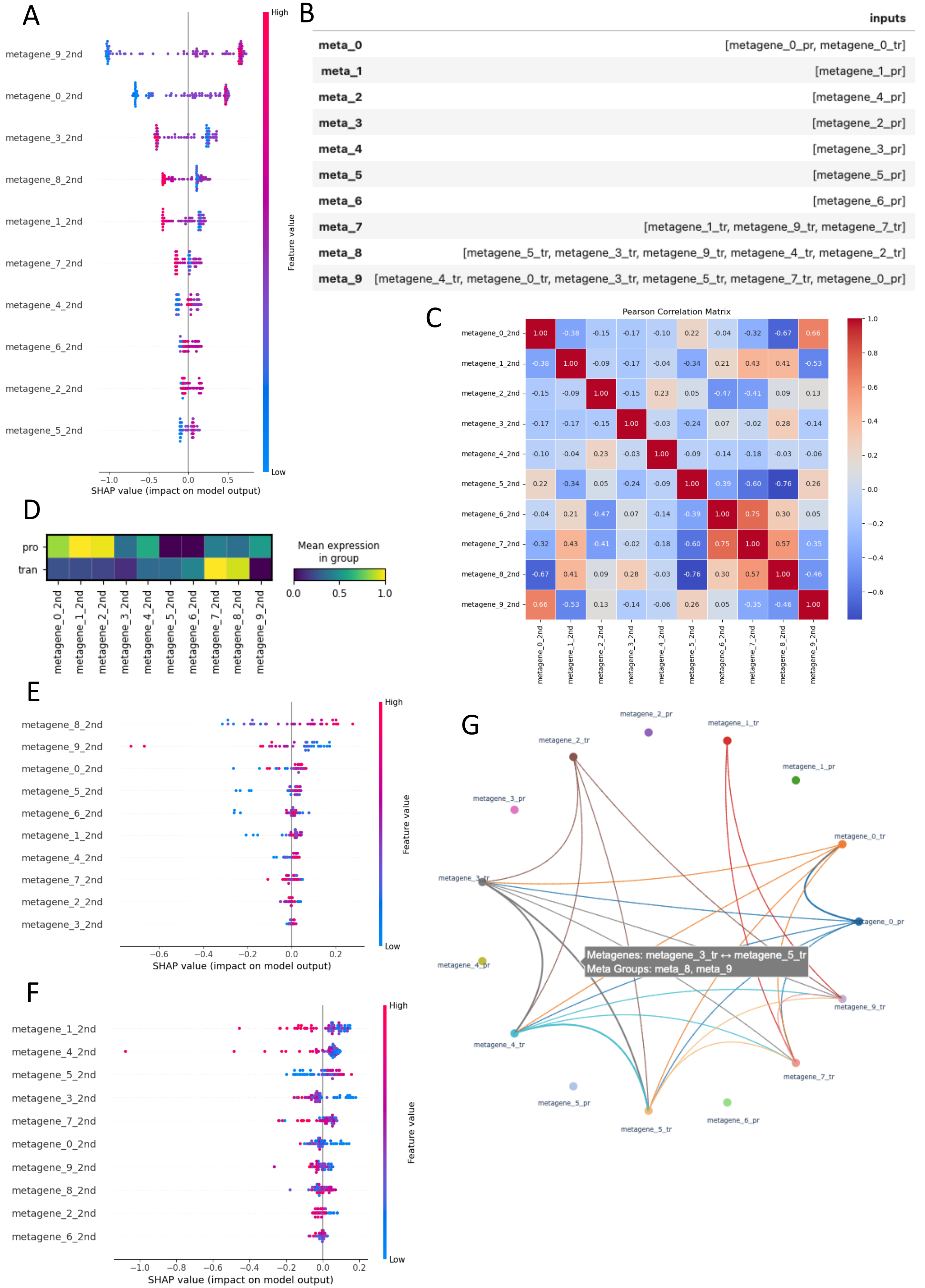


| | inputs |
|---|---|
| meta_0 | [metagene_0_pr, metagene_0_tr] |
| meta_1 | [metagene_1_pr] |
| meta_2 | [metagene_4_pr] |
| meta_3 | [metagene_2_pr] |
| meta_4 | [metagene_3_pr] |
| meta_5 | [metagene_5_pr] |
| meta_6 | [metagene_6_pr] |
| meta_7 | [metagene_1_tr, metagene_9_tr, metagene_7_tr] |
| meta_8 | [metagene_5_tr, metagene_3_tr, metagene_9_tr, metagene_4_tr, metagene_2_tr] |
| meta_9 | [metagene_4_tr, metagene_0_tr, metagene_3_tr, metagene_5_tr, metagene_7_tr, metagene_0_pr] |

| | metagene_0_2nd | metagene_1_2nd | metagene_2_2nd | metagene_3_2nd | metagene_4_2nd | metagene_5_2nd | metagene_6_2nd | metagene_7_2nd | metagene_8_2nd | metagene_9_2nd |
|---|---|---|---|---|---|---|---|---|---|---|
| metagene_0_2nd | 1.00 | -0.38 | -0.15 | -0.17 | -0.10 | 0.22 | -0.04 | -0.32 | -0.67 | 0.66 |
| metagene_1_2nd | -0.38 | 1.00 | -0.09 | -0.17 | -0.04 | -0.34 | 0.21 | 0.43 | 0.41 | -0.53 |
| metagene_2_2nd | -0.15 | -0.09 | 1.00 | -0.15 | 0.23 | 0.05 | -0.47 | -0.41 | 0.09 | 0.13 |
| metagene_3_2nd | -0.17 | -0.17 | -0.15 | 1.00 | -0.03 | -0.24 | 0.07 | -0.02 | 0.28 | -0.14 |
| metagene_4_2nd | -0.10 | -0.04 | 0.23 | -0.03 | 1.00 | -0.09 | -0.14 | -0.18 | -0.03 | -0.06 |
| metagene_5_2nd | 0.22 | -0.34 | 0.05 | -0.24 | -0.09 | 1.00 | -0.39 | -0.60 | -0.76 | 0.26 |
| metagene_6_2nd | -0.04 | 0.21 | -0.47 | 0.07 | -0.14 | -0.39 | 1.00 | 0.75 | 0.30 | 0.05 |
| metagene_7_2nd | -0.32 | 0.43 | -0.41 | -0.02 | -0.18 | -0.60 | 0.75 | 1.00 | 0.57 | -0.35 |
| metagene_8_2nd | -0.67 | 0.41 | 0.09 | 0.28 | -0.03 | -0.76 | 0.30 | 0.57 | 1.00 | -0.46 |
| metagene_9_2nd | 0.66 | -0.53 | 0.13 | -0.14 | -0.06 | 0.26 | 0.05 | -0.35 | -0.46 | 1.00 |

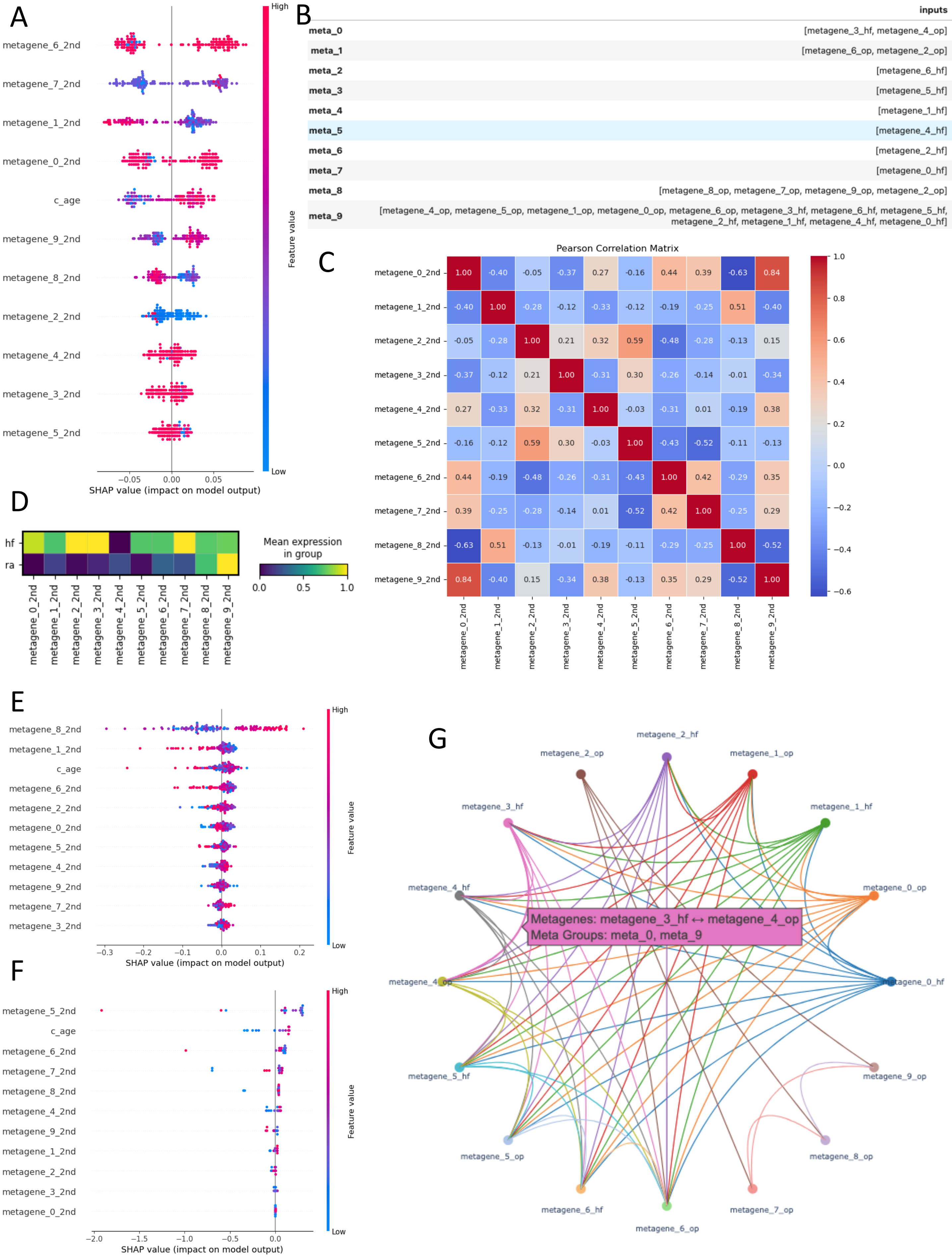
A
metagene_6_2nd
metagene_7_2nd
metagene_1_2nd
metagene_0_2nd
c_age
metagene_9_2nd
metagene_8_2nd
metagene_2_2nd
metagene_4_2nd
metagene_3_2nd
metagene_5_2nd
−0.05 0.00 0.05
SHAP value (impact on model output)
High
Feature value
Low
B
inputs
meta_0 [metagene_3_hf, metagene_4_op]
meta_1 [metagene_6_op, metagene_2_op]
meta_2 [metagene_6_hf]
meta_3 [metagene_5_hf]
meta_4 [metagene_1_hf]
meta_5 [metagene_4_hf]
meta_6 [metagene_2_hf]
meta_7 [metagene_0_hf]
meta_8 [metagene_8_op, metagene_7_op, metagene_9_op, metagene_2_op]
meta_9 [metagene_4_op, metagene_5_op, metagene_1_op, metagene_0_op, metagene_6_op, metagene_3_hf, metagene_6_hf, metagene_5_hf, metagene_2_hf, metagene_1_hf, metagene_4_hf, metagene_0_hf]
C
Pearson Correlation Matrix
metagene_0_2nd 1.00 -0.40 -0.05 -0.37 0.27 -0.16 0.44 0.39 -0.63 0.84
metagene_1_2nd -0.40 1.00 -0.28 -0.12 -0.33 -0.12 -0.19 -0.25 0.51 -0.40
metagene_2_2nd -0.05 -0.28 1.00 0.21 0.32 0.59 -0.48 -0.28 -0.13 0.15
metagene_3_2nd -0.37 -0.12 0.21 1.00 -0.31 0.30 -0.26 -0.14 -0.01 -0.34
metagene_4_2nd 0.27 -0.33 0.32 -0.31 1.00 -0.03 -0.31 0.01 -0.19 0.38
metagene_5_2nd -0.16 -0.12 0.59 0.30 -0.03 1.00 -0.43 -0.52 -0.11 -0.13
metagene_6_2nd 0.44 -0.19 -0.48 -0.26 -0.31 -0.43 1.00 0.42 -0.29 0.35
metagene_7_2nd 0.39 -0.25 -0.28 -0.14 0.01 -0.52 0.42 1.00 -0.25 0.29
metagene_8_2nd -0.63 0.51 -0.13 -0.01 -0.19 -0.11 -0.29 -0.25 1.00 -0.52
metagene_9_2nd 0.84 -0.40 0.15 -0.34 0.38 -0.13 0.35 0.29 -0.52 1.00
metagene_0_2nd metagene_1_2nd metagene_2_2nd metagene_3_2nd metagene_4_2nd metagene_5_2nd metagene_6_2nd metagene_7_2nd metagene_8_2nd metagene_9_2nd
1.0 0.8 0.6 0.4 0.2 0.0 −0.2 −0.4 −0.6
D
hf
ra
Mean expression in group
0.0 0.5 1.0
metagene_0_2nd metagene_1_2nd metagene_2_2nd metagene_3_2nd metagene_4_2nd metagene_5_2nd metagene_6_2nd metagene_7_2nd metagene_8_2nd metagene_9_2nd
E
metagene_8_2nd
metagene_1_2nd
c_age
metagene_6_2nd
metagene_2_2nd
metagene_0_2nd
metagene_5_2nd
metagene_4_2nd
metagene_9_2nd
metagene_7_2nd
metagene_3_2nd
−0.3 −0.2 −0.1 0.0 0.1 0.2
SHAP value (impact on model output)
High
Feature value
Low
F
metagene_5_2nd
c_age
metagene_6_2nd
metagene_7_2nd
metagene_8_2nd
metagene_4_2nd
metagene_9_2nd
metagene_1_2nd
metagene_2_2nd
metagene_3_2nd
metagene_0_2nd
−2.0 −1.5 −1.0 −0.5 0.0
SHAP value (impact on model output)
High
Feature value
Low
G
metagene_2_hf
metagene_2_op
metagene_1_op
metagene_3_hf
metagene_1_hf
metagene_4_hf
metagene_0_op
Metagenes: metagene_3_hf ↔ metagene_4_op
Meta Groups: meta_0, meta_9
metagene_4_op
metagene_0_hf
metagene_5_hf
metagene_9_op
metagene_5_op
metagene_8_op
metagene_6_hf
metagene_7_op
metagene_6_op

A

| | inputs |
|---|---|
| **meta_0** | [cell_type_Doublet, cell_type_Pre-ciliated, cell_type_Ionocyte, metagene_12_r4, metagene_10_r2, metagene_14_r2, metagene_11_r4, cell_type_Brush+PNEC, metagene_17_r1] |
| **meta_1** | [cell_type_Basal, metagene_12_r4, metagene_15_r1, cell_type_Pre-ciliated, metagene_9_r3, metagene_10_r4, metagene_8_r1, metagene_19_r2, metagene_12_r3, cell_type_Ionocyte, metagene_1_r2, metagene_10_r2, metagene_18_r4, metagene_16_r2, metagene_18_r2, metagene_12_r1, metagene_7_r3, metagene_3_r2, metagene_4_r4, cell_type_Ciliated] |
| **meta_2** | [cell_type_Hillock, metagene_13_r1, metagene_6_r2, metagene_10_r4, metagene_15_r2, cell_type_Pre-ciliated, metagene_19_r2, metagene_17_r3, metagene_9_r3, metagene_18_r4, metagene_16_r4, cell_type_Ionocyte] |
| **meta_3** | [metagene_11_r3, metagene_6_r1, metagene_13_r4, metagene_17_r2, metagene_15_r2, metagene_10_r4, cell_type_Brush+PNEC, metagene_14_r4, metagene_6_r4, metagene_14_r2, cell_type_Pre-ciliated, metagene_12_r2, metagene_9_r4, metagene_19_r2, metagene_9_r1, metagene_4_r3, metagene_5_r1, metagene_9_r2, metagene_7_r4, metagene_2_r2, metagene_16_r2, metagene_4_r1, metagene_12_r1, metagene_8_r4, metagene_1_r4] |
| **meta_4** | [cell_type_Secretory, metagene_15_r2] |

B

```
['cell_type_Doublet', 'cell_type_Pre-ciliated', 'cell_type_Ionocyte', 'cell_type_Brush+PNEC']
['cell_type_Basal', 'cell_type_Pre-ciliated', 'cell_type_Ionocyte', 'cell_type_Ciliated']
['cell_type_Hillock', 'cell_type_Pre-ciliated', 'cell_type_Ionocyte']
['cell_type_Brush+PNEC', 'cell_type_Pre-ciliated']
['cell_type_Secretory']
['cell_type_Cycling basal', 'cell_type_Ionocyte', 'cell_type_Pre-ciliated', 'cell_type_Ciliated']
['cell_type_Ciliated', 'cell_type_Brush+PNEC', 'cell_type_Cycling basal', 'cell_type_Ionocyte', 'cell_type_Basal']
['cell_type_Ionocyte', 'cell_type_Pre-ciliated']
['cell_type_Brush+PNEC', 'cell_type_Pre-ciliated']
['cell_type_Brush+PNEC', 'cell_type_Pre-ciliated', 'cell_type_Cycling basal']
['cell_type_Pre-ciliated', 'cell_type_Brush+PNEC']
['cell_type_Myoepithelial']
['cell_type_Ionocyte']
['cell_type_Brush+PNEC', 'cell_type_Cycling basal']
['cell_type_Brush+PNEC']
['cell_type_Brush+PNEC', 'cell_type_Myoepithelial', 'cell_type_Ionocyte']
['cell_type_Brush+PNEC']
['cell_type_Ionocyte', 'cell_type_Myoepithelial']
[]
['cell_type_Brush+PNEC', 'cell_type_Cycling basal', 'cell_type_Basal', 'cell_type_Doublet', 'cell_type_Hillock']
```

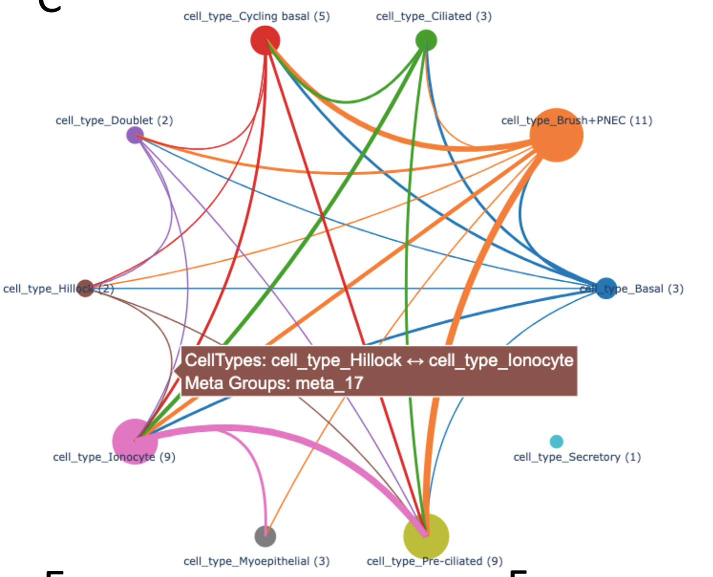


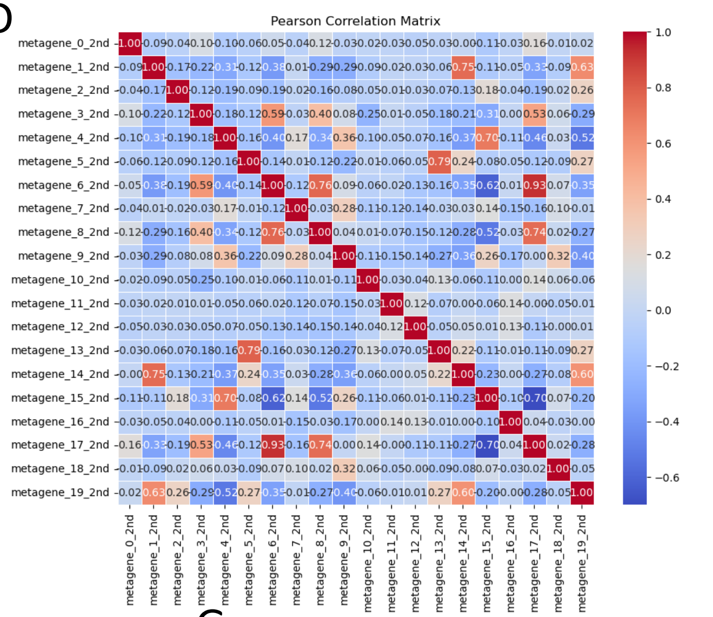


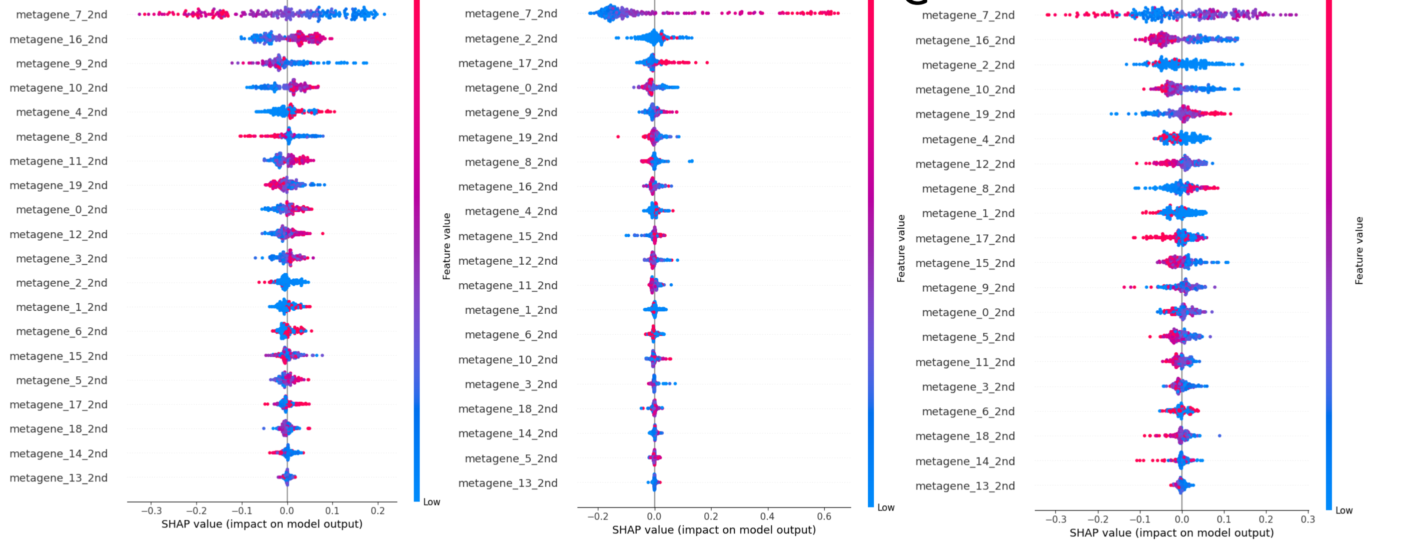

A

| | inputs |
|---|---|
| **meta_0** | [metagene_4_hum, metagene_9_hum, metagene_0_hum, metagene_4_plas, metagene_2_hum, metagene_3_plas, metagene_6_plas, metagene_8_plas] |
| **meta_1** | [metagene_6_hum] |
| **meta_2** | [metagene_3_hum] |
| **meta_3** | [metagene_2_hum, metagene_4_hum, metagene_5_hum, metagene_8_hum] |
| **meta_4** | [metagene_0_hum, metagene_1_plas, metagene_5_hum, metagene_2_plas, metagene_0_plas, metagene_4_hum, metagene_9_plas] |
| **meta_5** | [metagene_7_hum, metagene_5_hum, metagene_9_hum, metagene_4_hum] |
| **meta_6** | [metagene_1_hum, metagene_9_hum, metagene_8_hum, metagene_5_hum, metagene_4_hum] |

B

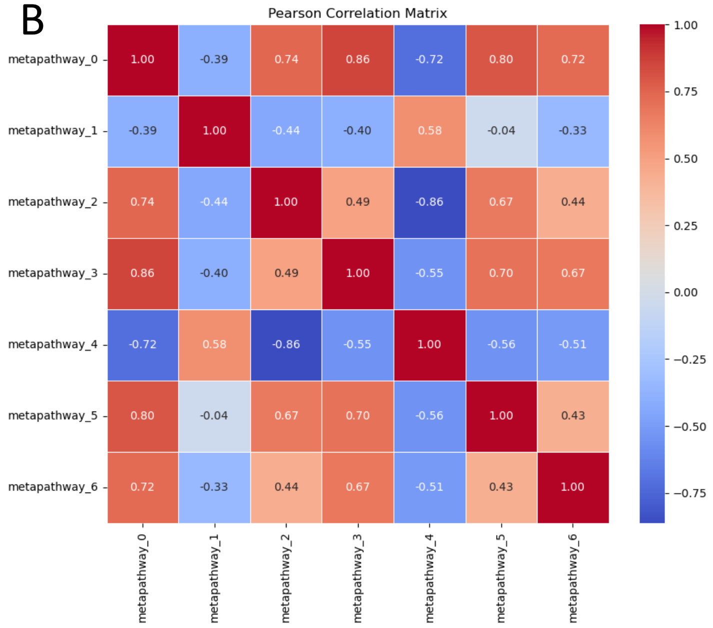


C

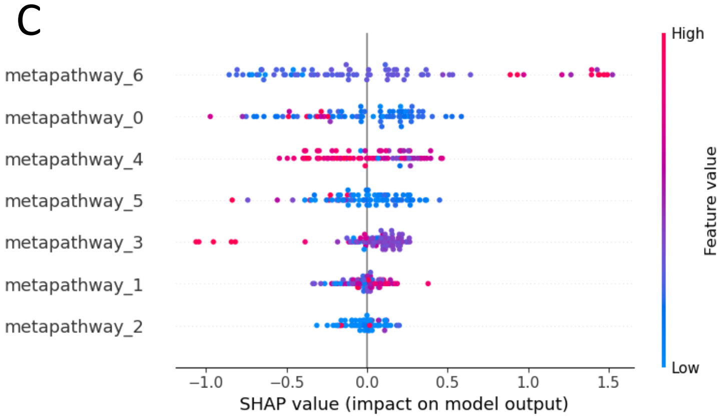

A

| | inputs |
|---|---|
| **meta_0** | [HSP90AA1, metagene_3_1st] |
| **meta_1** | [metagene_2_1st, metagene_0_1st, metagene_3_1st, metagene_13_1st] |
| **meta_2** | [metagene_1_1st, metagene_0_1st, metagene_8_1st, metagene_7_1st, metagene_4_1st, metagene_11_1st] |
| **meta_3** | [metagene_19_1st, metagene_17_1st, metagene_16_1st, metagene_7_1st, metagene_5_1st, metagene_0_1st, metagene_14_1st] |
| **meta_4** | [metagene_5_1st, metagene_16_1st, metagene_9_1st, metagene_12_1st, metagene_18_1st, metagene_1_1st, metagene_4_1st, metagene_11_1st, metagene_2_1st, metagene_19_1st] |
| **meta_5** | [metagene_18_1st, metagene_17_1st, metagene_5_1st, metagene_7_1st] |
| **meta_6** | [metagene_15_1st, metagene_14_1st, metagene_4_1st, metagene_6_1st, metagene_0_1st, metagene_18_1st] |

B

| | inputs |
|---|---|
| **meta_0** | [metagene_8_1st, metagene_1_1st, metagene_14_1st, metagene_10_1st, metagene_9_1st, metagene_18_1st, metagene_6_1st, metagene_0_1st, metagene_2_1st, metagene_15_1st] |
| **meta_1** | [HSP90AA1, metagene_7_1st] |
| **meta_2** | [metagene_2_1st, metagene_0_1st, metagene_16_1st, metagene_7_1st, metagene_5_1st] |
| **meta_3** | [metagene_19_1st, metagene_4_1st, metagene_9_1st, metagene_16_1st] |
| **meta_4** | [metagene_15_1st, metagene_16_1st, metagene_6_1st, metagene_17_1st, metagene_14_1st] |
| **meta_5** | [metagene_18_1st, metagene_14_1st, metagene_17_1st, metagene_16_1st, metagene_8_1st, metagene_5_1st, metagene_4_1st] |
| **meta_6** | [metagene_10_1st, metagene_16_1st, metagene_13_1st, metagene_9_1st, metagene_11_1st, metagene_14_1st, metagene_12_1st, metagene_7_1st, metagene_8_1st, metagene_5_1st] |

C

| | inputs |
|---|---|
| **meta_0** | [metagene_1_1st, metagene_12_1st, metagene_0_1st, metagene_10_1st, metagene_13_1st, metagene_16_1st, metagene_4_1st, metagene_7_1st, metagene_8_1st] |
| **meta_1** | [metagene_2_1st, metagene_11_1st, metagene_0_1st, metagene_7_1st, metagene_15_1st, metagene_9_1st] |
| **meta_2** | [metagene_19_1st, metagene_4_1st, metagene_16_1st, metagene_9_1st, metagene_12_1st, metagene_17_1st, metagene_8_1st, metagene_15_1st, metagene_7_1st] |
| **meta_3** | [metagene_18_1st, metagene_17_1st, metagene_0_1st, metagene_8_1st, metagene_9_1st, metagene_16_1st] |
| **meta_4** | [metagene_15_1st, metagene_16_1st, metagene_3_1st, metagene_6_1st, metagene_0_1st, metagene_19_1st, metagene_2_1st] |
| **meta_5** | [HSP90AA1] |
| **meta_6** | [metagene_8_1st, metagene_9_1st, metagene_14_1st, metagene_13_1st, metagene_3_1st, metagene_1_1st, metagene_11_1st, metagene_15_1st, metagene_19_1st] |

D

| Name | PValue |
|---|---|
| REACTOME_INTERFERON_SIGNALING | 1.180447e-46 |
| REACTOME_INTERFERON_ALPHA_BETA_SIGNALING | 9.281572e-46 |
| REACTOME_INTERFERON_GAMMA_SIGNALING | 8.758663e-32 |
| WP_IMMUNE_RESPONSE_TO_TUBERCULOSIS | 7.176071e-17 |
| WP_TYPE_II_INTERFERON_SIGNALING | 1.766636e-16 |
| WP_TYPE_II_INTERFERON_SIGNALING_IFNG | 3.867624e-13 |

E

| Name | PValue |
|---|---|
| KEGG_RIBOSOME | 3.837953e-70 |
| REACTOME_EUKARYOTIC_TRANSLATION_ELONGATION | 4.738412e-69 |
| REACTOME_SRP_DEPENDENT_COTRANSLATIONAL_PROTEIN_TARGETING_TO_MEMBRANE | 9.042303e-65 |
| KEGG_MEDICUS_REFERENCE_TRANSLATION_INITIATION | 1.956196e-64 |
| REACTOME_RESPONSE_OF_EIF2AK4_GCN2_TO_AMINO_ACID_DEFICIENCY | 6.309573e-64 |
| REACTOME_TRANSLATION | 7.070051e-64 |
| WP_CYTOPLASMIC_RIBOSOMAL_PROTEINS | 3.775265e-61 |
| REACTOME_NONSENSE_MEDIATED_DECAY_NMD_INDEPENDENT_OF_THE_EXON_JUNCTION_COMPLEX_EJC | 1.401779e-59 |
| REACTOME_TRANSLATION | 1.783624e-59 |
| REACTOME_NONSENSE_MEDIATED_DECAY_NMD | 5.048177e-59 |

F

| Name | PValue |
|---|---|
| BIOCARTA_AHSP_PATHWAY | 6.410068e-08 |
| REACTOME_HEME_BIOSYNTHESIS | 1.245165e-07 |
| REACTOME_ANTIGEN_PROCESSING_UBIQUITINATION_PROTEASOME_DEGRADATION | 8.878838e-07 |
| REACTOME_ANTIGEN_PROCESSING_UBIQUITINATION_PROTEASOME_DEGRADATION | 1.602668e-06 |
| REACTOME_METABOLISM_OF_PORPHYRINS | 1.855761e-06 |
| REACTOME_HEME_BIOSYNTHESIS | 1.893984e-06 |
| REACTOME_CELLULAR_RESPONSE_TO_CHEMICAL_STRESS | 1.985276e-06 |
| REACTOME_CELLULAR_RESPONSE_TO_CHEMICAL_STRESS | 2.139330e-06 |
| REACTOME_L1CAM_INTERACTIONS | 2.236713e-06 |

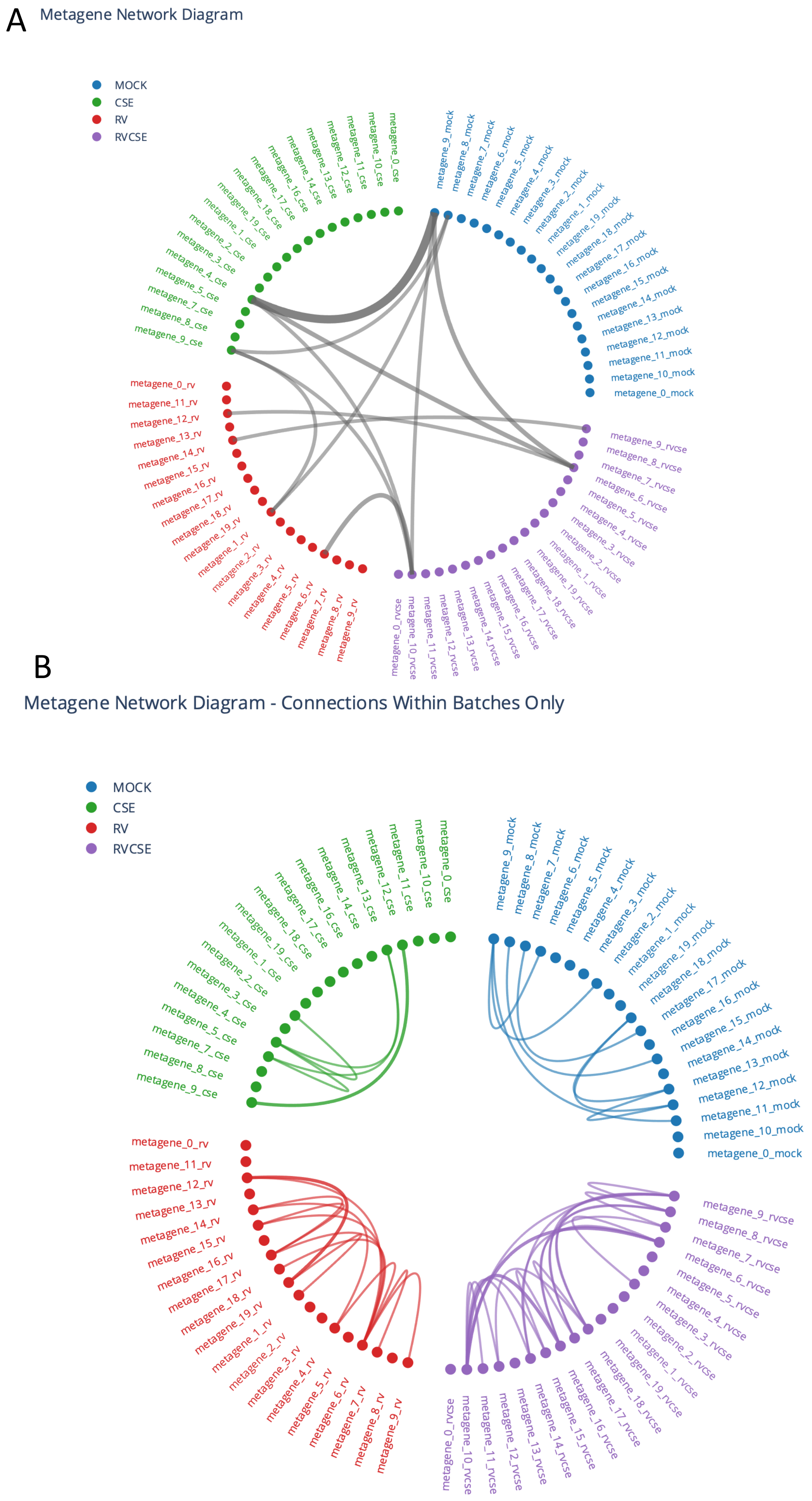
A
Metagene Network Diagram
MOCK
CSE
RV
RVCSE
metagene_0_cse
metagene_10_cse
metagene_11_cse
metagene_12_cse
metagene_13_cse
metagene_14_cse
metagene_16_cse
metagene_17_cse
metagene_18_cse
metagene_19_cse
metagene_1_cse
metagene_2_cse
metagene_3_cse
metagene_4_cse
metagene_5_cse
metagene_7_cse
metagene_8_cse
metagene_9_cse
metagene_9_mock
metagene_8_mock
metagene_7_mock
metagene_6_mock
metagene_5_mock
metagene_4_mock
metagene_3_mock
metagene_2_mock
metagene_1_mock
metagene_19_mock
metagene_18_mock
metagene_17_mock
metagene_16_mock
metagene_15_mock
metagene_14_mock
metagene_13_mock
metagene_12_mock
metagene_11_mock
metagene_10_mock
metagene_0_mock
metagene_0_rv
metagene_11_rv
metagene_12_rv
metagene_13_rv
metagene_14_rv
metagene_15_rv
metagene_16_rv
metagene_17_rv
metagene_18_rv
metagene_19_rv
metagene_1_rv
metagene_2_rv
metagene_3_rv
metagene_4_rv
metagene_5_rv
metagene_6_rv
metagene_7_rv
metagene_8_rv
metagene_9_rv
metagene_9_rvcse
metagene_8_rvcse
metagene_7_rvcse
metagene_6_rvcse
metagene_5_rvcse
metagene_4_rvcse
metagene_3_rvcse
metagene_2_rvcse
metagene_1_rvcse
metagene_19_rvcse
metagene_18_rvcse
metagene_17_rvcse
metagene_16_rvcse
metagene_15_rvcse
metagene_14_rvcse
metagene_13_rvcse
metagene_12_rvcse
metagene_11_rvcse
metagene_10_rvcse
metagene_0_rvcse
B
Metagene Network Diagram - Connections Within Batches Only
MOCK
CSE
RV
RVCSE
